# Partial *in vivo* reprogramming enables injury-free intestinal regeneration via autonomous Ptgs1 induction

**DOI:** 10.1101/2023.02.25.530001

**Authors:** Jumee Kim, Somi Kim, Seung-Yeon Lee, Beom-Ki Jo, Ji-Young Oh, Eun-Ji Kwon, Keun-Tae Kim, Anish Ashok Adpaikar, Eun-Jung Kim, Han-Sung Jung, Chang Pyo Hong, Jong Kyoung Kim, Bon Kyoung Koo, Hyuk-Jin Cha

## Abstract

Tissue regeneration after injury involves the dedifferentiation of somatic cells, a natural adaptive reprogramming process that leads to the emergence of injury-responsive cells with fetal-like characteristics in the intestinal epithelium. However, there is no direct evidence that adaptive reprogramming involves a shared molecular mechanism with direct cellular reprogramming. Here, we induced dedifferentiation of intestinal epithelial cells through forced partial reprogramming *in vivo* using “Yamanaka factors” (Oct4, Sox2, Klf4, and c-Myc: OSKM). The OSKM-induced dedifferentiation showed similar molecular features of intestinal regeneration, including a rapid transition from homeostatic cell types to injury-responsive-like cell types. These injury-responsive-like cells, sharing a gene signature of revival stem cells and atrophy-induced villus epithelial cells, actively assisted tissue regeneration following ionizing radiation-induced acute tissue damage. In contrast to normal intestinal regeneration, which involves epi-mesenchymal crosstalk through induction of *Ptgs2* (encoding Cox2) upon injury, the OSKM expression promotes the autonomous production of prostaglandin E2 via epithelial *Ptgs1* (encoding Cox1) expression. These results indicate that prostaglandin synthesis is a common mechanism for intestine epithelial regeneration but involves a different enzyme (*Ptgs1* for Cox1) when partial reprogramming is directly applied to the intestinal epithelium.

## Introduction

In the intestinal epithelium, Lgr5+ crypt base columnar cells (CBC cells) play the workhorse stem cell role by constantly producing progenitors and differentiated cells during homeostasis (Barker, 2014; Tan et al., 2021). As they are constantly undergoing cell division, Lgr5+ CBC cells are highly sensitive to genotoxic stresses such as ionizing radiation (IR). Upon IR-induced crypt damage, the cycling CBC cells are completely lost, yet the intestinal epithelium is still regenerated and can restore the Lgr5+ CBC cells (Tian et al., 2011). Various reserve stem cells and injury-responsive cells have been proposed, including secretory progenitors (Jadhav et al., 2017; Murata et al., 2020; van Es et al., 2012), label-retaining +4 cells (Higa et al., 2022; Tian *et al*., 2011), and revival stem cells (Ayyaz et al., 2019). The consensus among these studies is that the intestinal epithelium possesses various populations of cells with varying degrees of plasticity, which can be recruited rapidly for the optimal healing process.

With the advent of single-cell RNA sequencing (scRNA-seq) technology, injury-responsive cells in the intestinal epithelium have been characterized, leading to the discovery of revival stem cells (revSCs) (Ayyaz *et al*., 2019) and atrophy-induced villus epithelial cells (aVECs) (Ohara et al., 2022). They appear in different parts of the intestinal units, *i*.*e*., crypts and villi, but share molecular characteristics: cellular plasticity, a fetal-like gene expression profile and yes-associated protein (YAP) activation (Ayyaz *et al*., 2019; Ohara *et al*., 2022). Due to the induction of a fetal gene expression program, the intestinal epithelial injury response is believed to involve dedifferentiation and induction of plasticity through natural adaptive reprogramming *in vivo* (Jessen et al., 2015).

Cellular plasticity and dedifferentiation of somatic cells up to pluripotency can be induced by prolonged expression of the ‘Yamanaka factors’, Oct4, Sox2, Klf4, and c-Myc (hereafter OSKM) (Takahashi and Yamanaka, 2006), through epigenetic reprogramming such as progressive and conserved global erasure of DNA methylation (De Carvalho et al., 2010). OSKM-mediated cellular reprogramming is the gold standard method for the establishment of induced pluripotent stem cells (iPSCs) and has been widely used for various applications (Hartman et al., 2020). More recently, several studies demonstrated that forced partial reprogramming with OSKM enables rejuvenation or fetal-like dedifferentiation of cells in the eye, muscle, heart, and liver by inducing youthful DNA methylation patterns and transcriptomes (Lu et al., 2020) and activating muscle stem cells and cardiomyocytes (Chen et al., 2021b; Hishida et al., 2022; Wang et al., 2021). However, no mechanistic insight has been revealed to account for the regenerative effect of partial reprogramming in various tissues.

Here, we hypothesized that the natural dedifferentiation and acquisition of plasticity (*i*.*e*., natural adaptive reprogramming) that are induced by intestinal tissue injury share common features of forced partial reprogramming, as they also induce a fetal-like phenotype with improved regeneration capacity. To test this hypothesis, we induced dedifferentiation of intestinal epithelial cells by forced partial reprogramming through the induction of OSKM *in vivo*. Through extensive scRNA-seq analysis, we found that OSKM drives the induction of dedifferentiation, resulting in injury-responsive-like cells (*e*.*g*., revSC and aVEC-like cells) similar to natural adaptive reprogramming process. These OSKM-induced dedifferentiation facilitated intestinal injury repair. Both ‘Injury-induced’ and ‘OSKM induced’ differentiation showed prostaglandin synthesis as a common underlying mechanism responsible for the regeneration. However, the OSKM-induced dedifferentiation involved the ectopic epithelial expression of prostaglandin-endoperoxide synthase 1 (*Ptgs1)*, not via mesenchymal activation of *Ptgs2* as for natural regeneration (Roulis et al., 2020). We found that *Ptgs1*-mediated epithelial prostaglandin production is the key event for *in vivo* partial reprogramming, which directly renders intestinal epithelial cells to adopt fetal characteristics without involving any mesenchymal assistance.

## Results

### Dedifferentiation of intestinal epithelium by partial reprogramming

Unlike chronic OSKM expression, which induces multiple teratomas in mice (Abad et al., 2013), transient OSKM induction (or partial reprogramming) improves aging phenotypes without tumor formation (Browder et al., 2022a; Ocampo et al., 2016). We induced OSKM in the intestine (Figure S1A) by doxycycline (Dox) administration for 4 days with neither tumor formation (Figure S1B) nor weight loss (Figure S1C) in the ‘reprogrammable mouse’ model (inducible OSKM: iOSKM) (Stadtfeld et al., 2010) (Figure 1A). Oct4 and Sox2 proteins, which are rarely expressed in the intestine, were induced throughout the intestinal epithelium, corresponding to clear 2A peptide expression after Dox challenge (Figure 1B). Upon transient OSKM expression, differentiated cells such as Paneth cells [(PCs; positive for lysozyme staining (Lyz)] and goblet cells [GCs; positive for Alcian blue-PAS staining (AB-PAS)] appeared diminished (Figure 1C), which was consistent with repression of marker genes for PCs (*i*.*e*., *Lyz*) and GCs (*i*.*e*., *Muc2*) (Figure 1D). Unlike the differentiated intestinal epithelial cells, the number of CBC stem cells, marked by olfactomedin-4 (Olfm4) (van der Flier et al., 2009), was not altered (Figure 1E), suggesting that transient OSKM induction leads to dedifferentiation of intestinal epithelial cells with minimal effects on CBC stem cells. Interestingly, the Dox-challenged intestine exhibited a hyperplastic morphology with lengthened crypt region (Figure 1F) and increased numbers of proliferating cells (Figures 1G and S1D).

**Figure 1.**
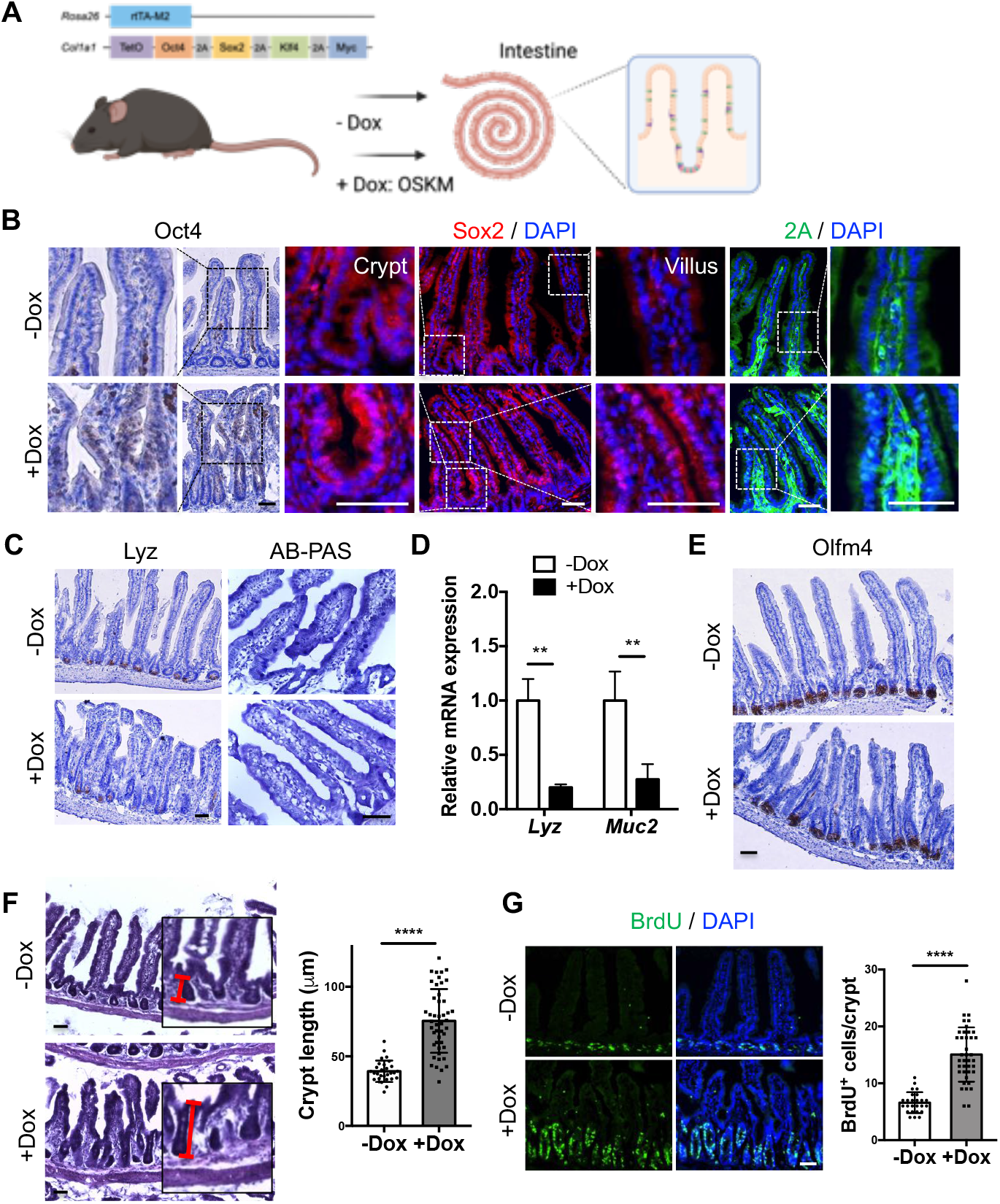
Induced dedifferentiation of the intestine by partial reprogramming. (A) Experimental scheme on the intestine of Dox-inducible OSKM (iOSKM) mouse model. (B) Immunohistochemistry (IHC) of Oct4 and Immunofluorescence (IF) of Sox2 and 2A peptide in the intestine of iOSKM mice. Samples were analysed 4 days after Dox administration. Hematoxylin or DAPI for nuclear staining. (C) Staining for Paneth cell (IHC of Lysozyme) and goblet cell (Alcian blue-PAS staining) in the intestine of iOSKM mice. (D) Relative mRNA expressions of *Lysozyme* (*Lyz* for Paneth cell) and *Mucin 2* (*Muc2* for goblet cell) in intestinal epithelial cells of iOSKM mice. n = 6; 3 mice□x□2 technical replicates. (E) IHC of Olfm4 in the intestine of iOSKM mice. (F) H&E histology in the intestine of iOSKM mice. Red line indicates the length of crypt (left). Quantification of crypt length (-Dox, n = 29; +Dox, n = 46 in total 3 mice of each condition) (right). (G) IF of BrdU in the intestine (left) and quantification of BrdU-positive cells per crypt (-Dox, n = 25; +Dox, n = 39 in total 3 mice of each condition) (right). BrdU was administered 3 hr prior to the sacrifice. Data represent the mean with SD (n = 10; 5 mice□x□2 technical replicates). Data represent the mean with SD. Student’s t-test: p <0.01(**), p <0.0001(****). Scale bar = 50μm.

### Cellular and molecular alterations induced by partial reprogramming

To characterize the cellular and molecular alterations induced by forced partial reprogramming, we performed scRNA-seq on intestinal epithelial cells with or without Dox administration (Figure 2A). Based on our computational pipeline for quality control (QC) and batch correction, we generated a partially reprogrammed intestine atlas of 17,000 QC-positive epithelial cells, with an average of 2,473 genes and 12,277 unique molecular identifiers (UMIs) per cell (Figure S1E). We applied uniform manifold approximation and projection (UMAP) for visualization and a graph-based clustering algorithm to identify 18 cell clusters (Figure S1F). Based on the expression of canonical marker genes, we identified 7 canonical cell types including crypt-base columnar cells (CBC), transit-amplifying cells (TA), enterocytes (EC), enteroendocrine cells (EE), goblet cells (GC), Paneth cells (PC) and tuft cells (TC), as well as two distinct Dox-induced subsets (DC1 and DC2) that were manifest after Dox treatment (Figures 2B, S1G, and S1H). Interestingly, DC1 did not express any canonical intestinal epithelial cell markers, while DC2 partially expressed EC-related marker genes (Figure S1G).

**Figure 2.**
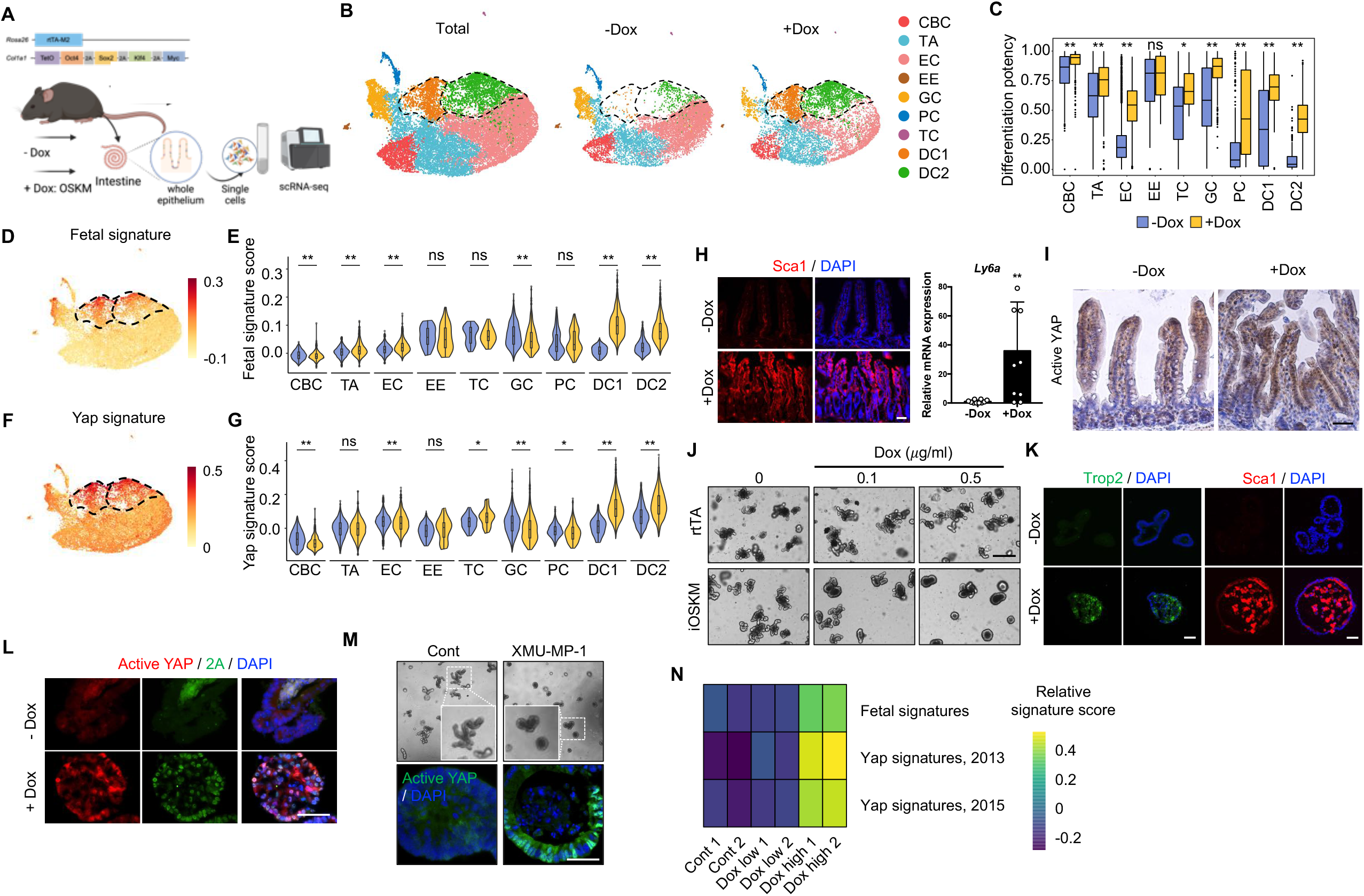
Concurrent fetal gene transition YAP activation and by partial reprogramming. (A) Experimental scheme for single cell RNA-seq using intestine of iOSKM mouse in Dox condition (B) UMAP plots of scRNA-seq from OSKM-induced mouse intestinal epithelium in –Dox and +Dox conditions (Total, left), -Dox (control, middle), and +Dox (Dox-treated, right). (C) Bar plot indicating differentiation potency inferred by CytoTRACE per each condition in different cell types. (D-E) UMAP plot (D) and violin plot (E) showing gene module score of 1,275 fetal signature genes. (F-G) UMAP plot (F) and violin plot (G) showing gene module score of 398 yap signature genes. (H) IF of Sca1 in the intestine (left). Relative mRNA expression of *Ly6a* (encoding Sca1) in intestinal epithelial cells of iOSKM mice after Dox treatment (right). DAPI for nuclear staining. (I) IHC of Active YAP in the intestine of iOSKM mice 4 days after Dox administration. (J) Microscopic images of Dox-treated intestinal organoids from rtTA mouse or iOSKM mouse. (K) IF of fetal markers, Trop2 and Sca1, in iOSKM intestinal organoids with Dox treatment for 3 days. DAPI for nuclear staining. (L) IF of Active YAP and 2A peptide in iOSKM intestinal organoids. DAPI for nuclear staining. (M) Microscopic images (top) and IF of Active YAP (bottom) in XMU-MP-1 (Mst1/2 inhibitor)-treated intestinal organoids. (N) Gene enrichment of YAP or Fetal signatures in intestinal organoids. Data represent the mean with SD. Student’s t-test: p < 0.05(*), p <0.01(**), ns, not significant. Scale bar = 50μm (H, I, K, L and M), 500μm (J).

We next examined the molecular alterations induced by forced partial reprogramming for each cell type. By measuring the extent of stemness or dedifferentiation potential, we found that most cell types except EE and TC showed significantly elevated stemness scores upon Dox treatment (Figures 2C and S1I). In addition, the expression of canonical marker genes for EE, GC, PC, and TC was decreased upon Dox treatment compared to their corresponding cell types (Figure S1J), in congruence with the observations in Figures 1C and D. The induction of a fetal gene signature was most evident in DC1 and DC2 clusters (Figures 2D and E), along with an enriched Yap signature (Figures 2F and G). It is noteworthy that the acquisition of fetal phenotypes during injury repair in the intestine also occurs in a Yap-dependent manner (Yui et al., 2018a). As predicted, we also observed induction of Sca1 (Figure 2H) and Trop2 (Figure S1K), encoded by *Ly6a* and *TACSTD2*, typical fetal genes expressed in the intestine after injury (Mustata et al., 2013; Nusse et al., 2018; Yui *et al*., 2018a), along with a pronounced nuclear translocated active Yap (unphosphorylated Yap) signal (Si et al., 2017) throughout the epithelium upon Dox treatment (Figure 2I). These results suggest that forced partial reprogramming induces emergence of two distinctive populations with fetal-like characteristics and YAP activation (DC1 and DC2) as well as cellular plasticity of diverse cell types.

Next, to examine the effect of OSKM directly on the intestinal epithelium, we established an intestinal organoid model from the ‘reprogrammable mouse’ (iOSKM). Transient OSKM expression for 3 days (Figures S2A and B) was insufficient to induce *Nanog*, a marker for full reprogramming, even with 0.5 μg/ml of Dox (Figure S2C), confirming that reprogramming was only partial and pluripotency was not induced. Interestingly, there was a clear morphological change from budding organoids to cystic spheroids upon Dox treatment in a dose-dependent manner, compared to Dox-treated intestinal organoids from *Rosa26-rtTA* mouse as a control of effects of Dox or rtTA expression (Figures 2J and S2D and Movie S1). This cystic spheroid phenotype was gradually lost after Dox withdrawal (Figure S2E), showing the reversible nature of the reprogramming, similar to the later recovery stage of injury response *in vivo* (Sato et al., 2020). Consistent with the acquisition of the ‘fetal-like gene signature’ by partial reprogramming of the intestine *in vivo* (Figures 2D and E), cystic spheroids resulting from OSKM induction resembled spheroids established from fetal mouse intestine (Figure S2F): they were positive for Sca1 and Trop2 (Figures 2K and S2G) and expressed the fetal genes *Ly6a* (encoding Sca1), *Anxa1*, and *Tacstd2* (encoding Trop2) in a dose-dependent manner (Figure S2H). This was clearly correlated with active YAP signal (Figure 2L) and expression of the Yap downstream target genes *Ccn2* and *Tead4* (Figure S2I). It is also noteworthy that Yap activation with XMU-MP-1, Mst1/2 inhibitor, induced cystic spheroid formation with clear active Yap signal (Figure 2M). Consistently, bulk RNA-seq using organoids also showed that OSKM induction (Figures S2K and S2J) led to enrichment of the fetal and Yap signatures (Figure 2N).

### Generation of two distinct injury-responsive-like cells by partial reprogramming

Previous studies demonstrated that two distinct populations, revSCs and aVECs, appear via (de)differentiation in the crypt and villi, respectively, upon injury and repair the damage (Ayyaz *et al*., 2019; Ohara *et al*., 2022). To examine the characteristics of DC1 and DC2, the newly formed clusters with fetal gene signatures we identified after partial reprogramming (Figures 2B and D), the transcriptome profiles of DC1 and DC2 were compared with those from revSCs induced by crypt damage with IR (Figures 3A and S3A) and aVECs induced by villi atrophy (Figures 3A and S3B). Interestingly, the DC1 subset was characterized by high expression of revSC markers (87 genes including *Clu, Anxa1, Ccnd1*, and *Ly6a*) (Figures 3B, S3C, and S3D), while the DC2 subset was distinguished from DC1 by mature EC markers highly expressed at the top of villi (353 genes including *Ada, Fabp2, Apoa4, Apoa1*, and *Alpi*) (Figures 3B, S3C, and S3E). Among reported aVEC markers, *Clu* and *Anxa1* were mainly found in DC1 (Figure 3C), while *Itgb6, Plaur* (Figure 3C), *Cldn4, Lamc2* (Figure S3F) as well as *Msln* (Figure 3D), were expressed in both DC1 and DC2 likely because of existence of overlapping makers for revSC and aVEC (Figure S3G). These results suggest that DC1 and DC2 considerably share molecular features with injury-induced revSC and aVEC, respectively.

**Figure 3.**
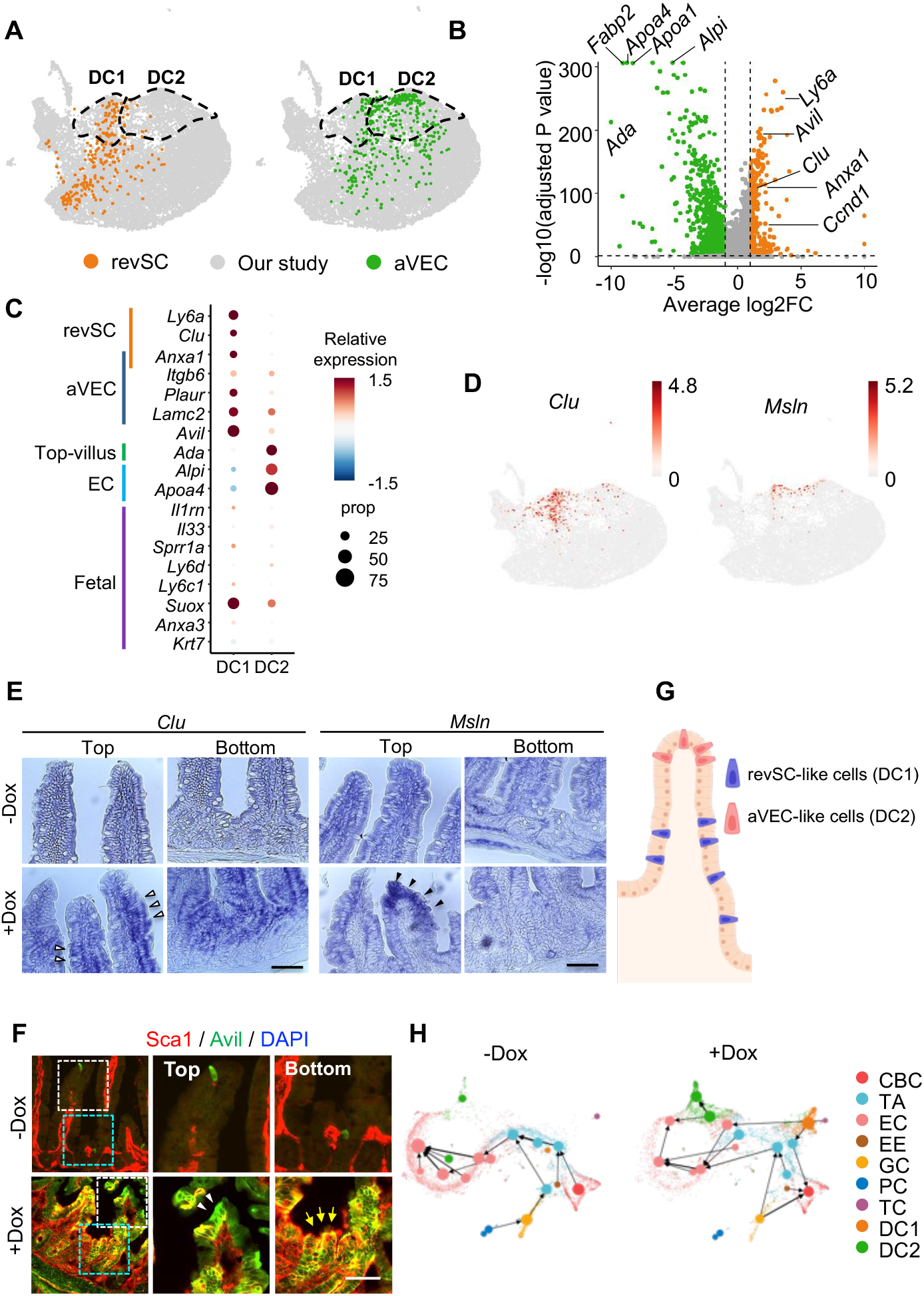
Generation of binary injury-responsive-like cells by partial reprogramming. (A) Projection of revSC (left) and aVEC (right) cells onto our scRNA-seq data. Cells in our study are indicated in gray. (B) Volcano plot showing differentially expressed genes (DEGs) between DC1 (right) and DC2 (left). (C) Dot plot for relative expression of marker genes of revSC, aVEC, top-villus, EC and fetal signature genes in DC1 and DC2. (D) UMAP plots showing gene expression of *Clu* and *Msln*. (E) In situ hybridization (ISH) of *Clu* and *Msln* in the intestine of iOSKM mice. white and black arrowheads indicate middle and top of the villi, respectively. White arrowhead indicates *Clu*^+^ revSC-like cells, while black arrowhead indicates *Msln*^+^ aVEC-like cells. (F) IF of Sca1 and Avil in the intestine of iOSKM mice. White arrowhead indicates Sca1^-^Avil^+^ aVEC-like cells, while yellow arrow indicates Sca1^+^Avli^+^ revSC-like cells. (G) Graphical presentation of spatial presence of revSC (DC1) or aVEC (DC2)- like cells (H) PAGA graphs showing velocity-directed arrows from cluster to cluster in –Dox (left) and +Dox (right). Scale bar = 50μm.

Subsequent RNA in situ hybridization revealed that *Clu*, exclusively expressed in DC1 (Figure 3D), was mainly elevated in the middle to the bottom of the villi as well as in the crypts (Figure 3E, left panels), while *Msln* (Figure 3E, right panels), expressed in DC1 and DC2 (Figure 3D), was elevated at the top of the villi. It was consistent with the result that top villus genes, including *Ada* (Moor et al., 2018) were enriched in DC2, not DC1 (Figures S3E and H). The discrete positions of DC1 and DC2 (Figure 3E) were also confirmed by immunostaining with Sca1 (encoded by *Ly6a* expressed in DC1) and Avil (expressed in both DC1 and DC2) (Figure 3F and S3I). The Sca1^-^Avil^+^ population resides at the top of the villi, representing DC2, while double-positive cells, representing DC1, are mostly found in the bottom villi as well as crypts (Figures 3F and G). As expected, our trajectory analyses positioned DC1 and DC2 close to EC and TA/CBC (Figures 3H and S3J). Accordingly, we defined the DC1 and DC2, populations produced by partial OSKM induction even in the absence of physical injury, as revSC-like and aVEC-like cells, respectively.

### Promotion of intestinal repair by partial reprogramming

The production of OSKM-induced injury-responsive-like cells led us to hypothesize that partial reprogramming may assist intestinal regeneration after injury. Since Oct4 expression was evident by 3 days after Dox administration (Figure S4A), ‘reprogrammable mice’ were pre-treated with Dox 2 days before IR. the intestines were harvested on day 2 and day 4 after IR-induced injury to examine the effect of partial reprogramming on acute tissue damage and the subsequent repair process (Figure 4A). On day 2 after IR, clear depletion of Olfm4 and Ki67 signals in the crypt was observed regardless of Dox treatment (Figure 4B), implying that similar crypt damage had occurred. In sharp contrast, on day 4, Dox-treated mice exhibited complete epithelial repair, along with re-expression of Olfm4 and Ki67 in the crypt and the TA zone, while mice without Dox treatment did not (Figures 4C and S4B). This accelerated repair induced by OSKM expression was associated with a significant increase in proliferating cells in the TA zone (pulse-chased with BrdU incorporation) (Figure 4D), Trop2 positive fetal-like cells (Figure S4C) and robust Sca1 expression throughout the entire villi (Figure 4E) at 4 days after IR, along with increased active Yap signal (Figure 4F). Similarly, in intestinal organoids, although Dox-treatment of intestinal organoids did not affect the levels of cell death observed at 4 days after IR exposure (Figure S4D), OSKM induction partially prevented the subsequent severe organoid shrinkage induced by IR (Figure S4E).

**Figure 4.**
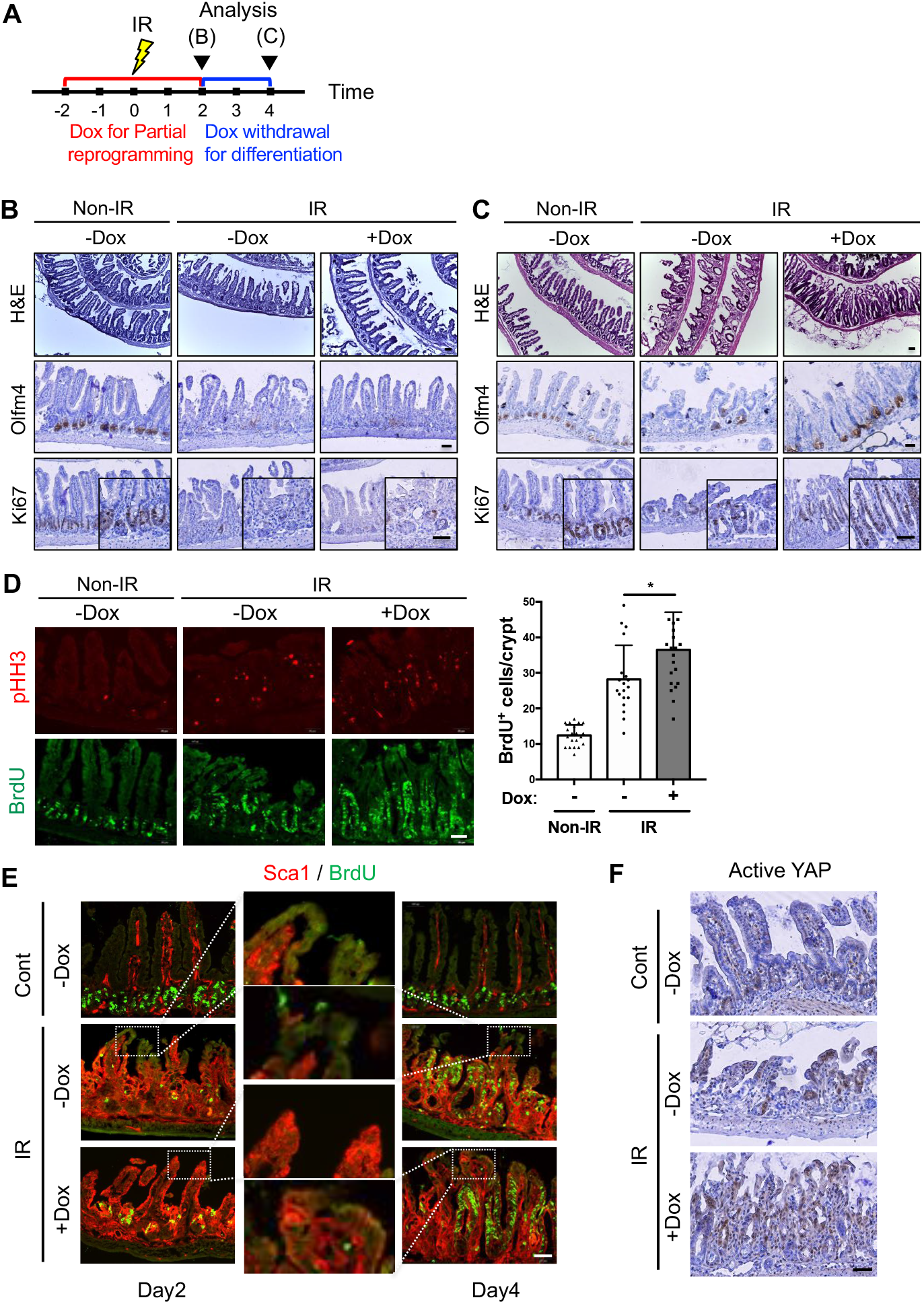
Facilitated intestinal repair after cryptic damage with IR by partial reprogramming. (A) Experimental scheme for 10Gy IR and Dox treatment in iOSKM mice. The day after IR was indicated. Dox was treated 2days before IR for 4 days, and then removed to produce differentiated cells from OSKM-induced dedifferentiated cells for regeneration. Analysis at 2dpi and 4dpi were showed in (B) and (C), respectively. (B and C) H&E histology and IHC of Olfm4 and Ki67 in the intestine of iOSKM mice after IR at 2dpi (days post-irradiation) (B) and at 4dpi (C). (D) IF of phospho-histone H3(pHH3) and BrdU in the intestine of iOSKM mice after IR at 4dpi (left) and quantification of BrdU+ cells per crypt (right). Data represent the mean with SD (-Dox and non-IR, n = 20; -Dox and IR, n = 19; +Dox and IR, n=22 in total 3 mice of each condition). p < 0.05(*). (E) IF of Sca1 and BrdU in the intestine of iOSKM mice after IR at 2dpi and 4dpi. Top of the villi were magnified. (F) IHC of Actve YAP in the intestine of iOSKM mice after IR at 4dpi. Data represent the mean with SD. Student’s t-test: p <0.05(*). Scale bar = 50μm.

### Partial reprogramming triggers epithelial prostaglandin E2 synthesis

Partial reprogramming produced injury-responsive-like cell populations (Figure 3) along with the acquisition of the fetal gene signature (Figure 2), similar to injury-induced dedifferentiation; thus, a common molecular mechanism was expected between partial reprogramming and injury-induced dedifferentiation (Figure 5A). To this end, we examined the commonly altered gene expression profiles of organoids after partial reprogramming and IR-mediated injury and found that a ‘Prostaglandin synthesis and Regulation’ signature was apparently most enriched (Figure 5A). The enrichment of the prostaglandin signature was commonly manifested in organoid from IR-mediated injury, fetal intestine, and partial reprogramming (Figure S5A). Consistently, significant induction of the prostaglandin signature was observed in DC1 and DC2 populations upon Dox treatment (Figure 5B). Of note, prostaglandins E2 (PGE2), naturally produced by cyclooxygenase (Cox) 1 and 2 (encoded by *Ptgs1* and *Ptgs2*, respectively), have been extensively studied in tissue regeneration (Cheng et al., 2021). It is also known that inhibition of PGE2 degradation promotes tissue regeneration of bone marrow, colon, and liver (Zhang et al., 2015) and aged muscle (Palla et al., 2021). While the natural epithelial regeneration of intestine is promoted by paracrine effect of PGE2 from mesenchymal Ptgs2 expression (Li et al., 2018; Miyoshi et al., 2017; Roulis *et al*., 2020), PGE2 production was observed from epithelium after OSKM induction but not from mesenchyme (Figure 5C).

**Figure 5.**
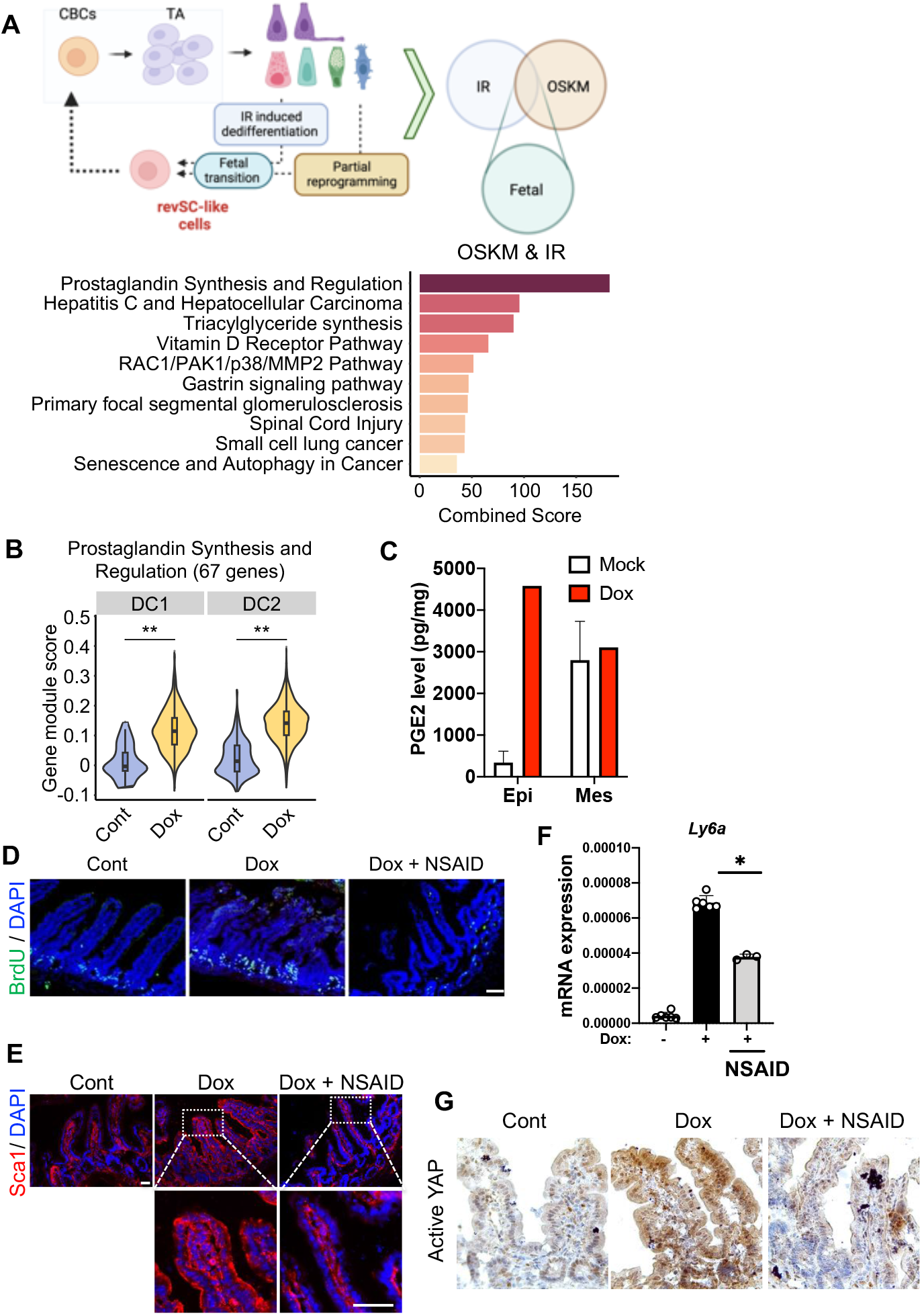
Prostaglandin synthesis for alterations induced by partial reprogramming. (A) A schematic model describing regeneration process in small intestine and common fetal transition (top), and a bar plot showing enrichR combined scores of commonly up-regulated pathways in OSKM-, IR-induced regenerating intestinal organoids (bottom). (B) A violin plot showing ‘Prostaglandin synthesis and Regulation’ signature score in cell types (C) PGE2 level (pg/mg) of mouse intestine in epithelium (Epi) and mesenchyme (Mes) with or without Dox treatment (D) IF of BrdU in the intestine of iOSKM mice, Cont (left), Dox (Dox-treated, middle), and Dox+NSAID (Dox and Dexibuprofen treated, right) (E) IF of fetal markers, Sca1, in iOSKM mice intestine treated with Dox and NSAID for 4days. DAPI for nuclear staining (F) mRNA expressions of *Ly6a* in intestinal epithelial cells of iOSKM mice. n = 6; 2mice□x□3 technical replicates. (G) IHC of Active YAP in the intestine of iOSKM mice after Dox and NSAID administration for 4 days

When treated nonsteroidal anti-inflammatory drug (NSAID), an inhibitor of Cox1 and Cox2, blocking the synthesis of PGE2, fetal-like transition by partial reprogramming was significantly compromised. The oral administration of dexibuprofen, a typical NSAID, reverted the effect of partial reprogramming, such as an increase of BrdU positive population in crypt (Figure 5D) and Sca1 positive population (Figure 5E), corresponding to the level of *Ly6a* expression (Figure 5F). Consistently, fetal-like phenotypes such as spheroid formation (Figure S5B) and Sca1 expression (Figures S5C and D) were reversed in the organoid model by NSAID treatment. We also observed that YAP activation in the intestine (Figure 5G) and intestinal organoid (Figure S5E) by partial reprogramming was also markedly attenuated by NSAID treatment.

### Cox1 activity is key for partial reprogramming-induced regeneration

The diminished YAP activity by NSAID (Figure 5G), prompted us to examine Cox1 and Cox2 more closely, since PGE2, of which production was manifested in intestine epithelium by OSKM induction (Figure 5C), is known to activate YAP through prostaglandin E–receptor 4 (EP4) (Kim et al., 2017). Quantitative RT-PCR data clearly revealed that *Ptgs1*, but not *Ptgs2*, was markedly induced by partial reprogramming (Figure 6A, left). scRNA-seq data also confirmed that *Ptgs1* expression was specifically upregulated in OSKM-induced clusters, DC1 and DC2 (Figure 6A, right). Cox1 expression, normally occurring in Dclk1 positive tuft cells (Gerbe et al., 2011), was shown to be induced predominantly in Dclk1 negative epithelial cells (Figure S6A), in both villus and crypt by OSKM induction unlike Cox2 (Figure 6B). The exclusive epithelial induction of *Ptgs1* by OSKM were manifested in intestine organoids compared to fetal organoids and organoids after IR-mediated injury (Figures S6B and C), along with expression of Sca1 and Avil (Figures S6D and E). It is noteworthy that *Ptgs1* expression normally remains constant, while *Ptgs2* is highly inducible under tissue damage and causes an inflammatory response (Vane et al., 1998). To determine the functionality of Cox1 induction (not Cox2) in the intestine epithelium by partial reprogramming, we utilized selective inhibitors of Cox1 and Cox2: SC-560, Cox1 inhibitor (iCox1); celecoxib, Cox2 inhibitor (iCox2). As predicted, inhibition of Cox1, but not Cox2 distinctively negated the effect of partial reprogramming, such as Sca1 expression, an increase of cryptic BrdU population (Figure 6C), formation of *Clu* positive population (Figure 6D), and YAP activation (Figure 6E). Accordingly, intestine regeneration and concurrent formation of Olfm4 positive cells at crypt by OSKM induction were markedly attenuated by treatment of Cox1 inhibitor but not Cox2 inhibitor (Figure 6F), which was corresponding to the level of BrdU positive population at crypt (Figure 6G). Taken together, autonomous epithelial production of PGE2 by Cox1 expression is a key molecular event for the induced dedifferentiation with partial reprogramming.

**Figure 6.**
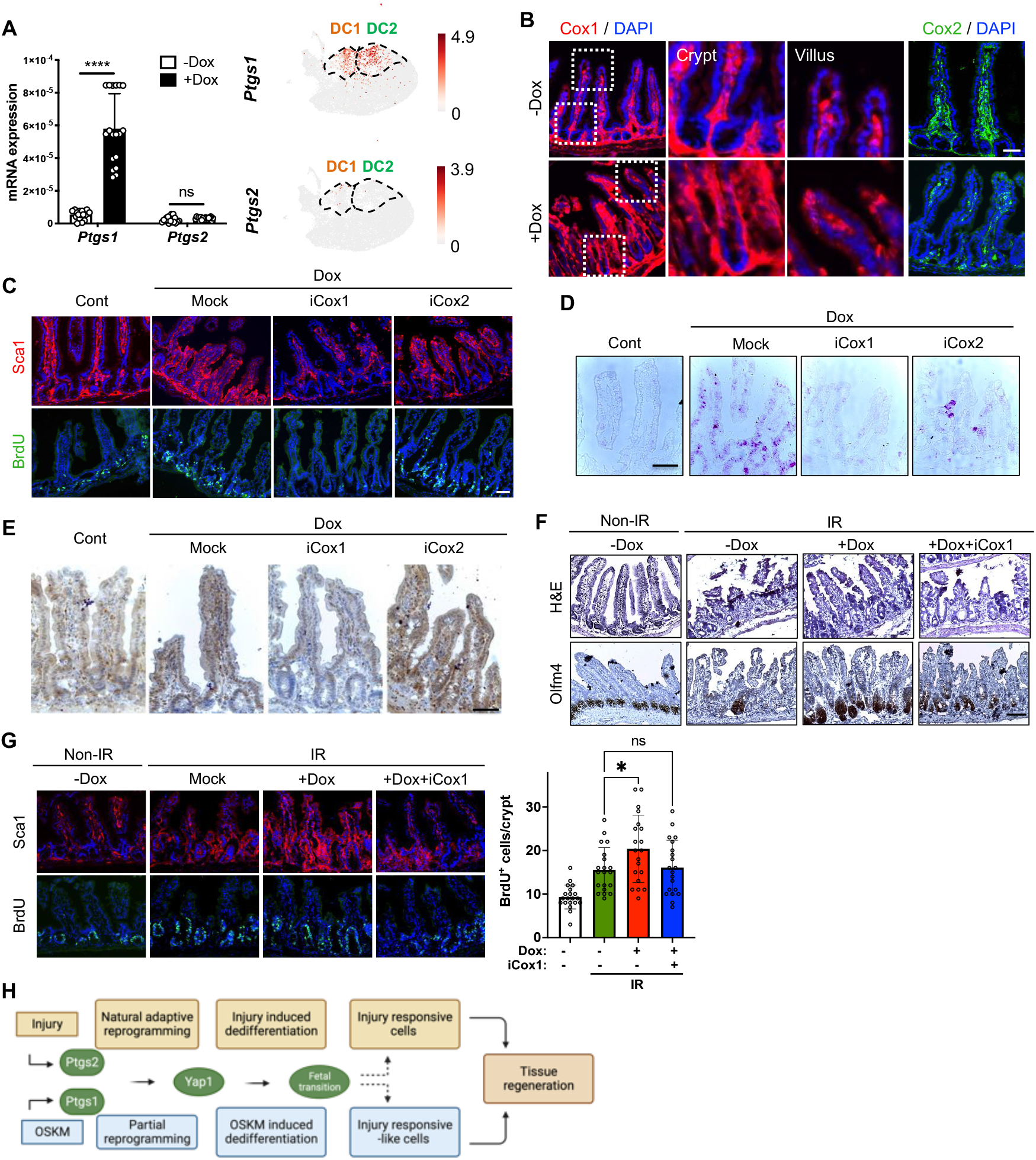
Induction of Cox1 in partial reprogramming-induced regeneration. (A) mRNA expressions of *Ly6a* in intestinal epithelial cells of iOSKM mice (right). n = 18; 6 mice□x□3 technical replicates (left). Projection of *Ptgs1* (right, top) and *Ptgs2* (right, bottom) cells over scRNA-seq data. Cells in our study indicated in gray. (B) IF of *Ptgs1*(left) and *Ptgs2*(right) in the intestine of iOSKM mice, 4 days after Dox administration. DAPI for nuclear staining. (C) IF of Sca1 and BrdU in iOSKM mice intestine in control, treated with Dox for 4 days, treated with Dox and Cox1 inhibitor (iCox1: SC-560) for 4 days and 2days respectively, and treated with Dox and Cox2 inhibitor (iCox2: Celecoxib) for 4 days and 2days respectively (from left to right). DAPI for nuclear staining. (D) In situ hybridization of *Clu* in the intestine of iOSKM mice (E) IHC of Active YAP in the intestine of iOSKM mice (F) H&E histology and IHC of Olfm4 in the intestine of iOSKM mice after IR and 4 day-Dox administration, 2-day Cox1 inhibition at 4dpi (G) IF of Sca1 and BrdU in the intestine of iOSKM mice (left), and quantification of BrdU-positive cells per crypt (n = 20 in total 3 mice for each condition) (right). (H) A schematic model for regeneration process induced by injury or OSKM

## Discussion

Despite multiple recent studies revealing regeneration induced by OSKM in diverse tissues (Chen *et al*., 2021b; Hishida *et al*., 2022; Lu *et al*., 2020; Wang *et al*., 2021), the phenotypic and mechanistic similarities between forced dedifferentiation and injury response-mediated (de)differentiation (Ayyaz *et al*., 2019; Ohara *et al*., 2022) remain undetermined. Here, based on scRNA-seq analysis, we demonstrated that the novel epithelial populations generated by OSKM-mediated partial reprogramming are comparable to injury-responsive subpopulations that are responsible for acute damage-repair in the crypts and villi, respectively. This observation was supported by the expression of specific markers such as *Clu, Msln, Sca1* and *Avil*, associated YAP activation and expression of fetal genes such as *Trop2* and *Sca1*. Induction of OSKM in organoids produced a fetal intestinal organoid phenotype (spheroids) with the expression of fetal gene signatures and activation of Yap. The induction of two distinct populations, DC1 and DC2, by OSKM expression mimics the generation of two injury-responsive cell types, revSC and aVEC, respectively. This means that a single cause (OSKM) can faithfully recapitulate the injury-mediated tissue dedifferentiation in the intestinal epithelium.

Notably, tissue damage induces Ptgs2 to promote PGE2 production not only for inflammatory response but also for damage repair (or regeneration) (Roulis *et al*., 2020) through ‘adaptive cellular response to produce wound repair cells’ (Miyoshi *et al*., 2017). In a similar manner, we showed that OSKM-mediated partial reprogramming, associated with fetal-transition by OSKM, was the result of an aberrant induction of *Ptgs1* and prostaglandin synthesis. These findings suggest that OSKM-induced (de)differentiation (via induction of epithelial *Ptgs1*) would share the same molecular mechanism (i.e., prostaglandin biosynthesis and subsequent Yap activation) as injury response-mediated (de)differentiation (through mesenchymal *Ptgs2* induction). It is noteworthy that there is emerging evidence that PGE2 is critical for tissue repair (or regeneration) of not only the intestine (Miyoshi *et al*., 2017; Roulis *et al*., 2020) but also the skeletal muscle (Ho et al., 2017), kidney (Chen et al., 2021a) and heart (Muraoka et al., 2019). Consistently, the inhibition of PGE2 degradation (Zhang *et al*., 2015) promotes both tissue regeneration and rejuvenation (Palla *et al*., 2021), as demonstrated in a mouse model by OSKM-induced partial reprogramming (Browder et al., 2022b; Hishida *et al*., 2022; Wang *et al*., 2021).

Unlike the formation of blastema by injury-induced dedifferentiation for appendage regeneration in amphibians or fish, only limited dedifferentiation upon injury occurs in mammals. Nevertheless, the genomic profile of blastema formation and the subsequent appendage is closely associated with prostaglandin biosynthesis and blocked by NSAID treatment, suggesting that prostaglandin synthesis is evolutionarily conserved for injury-induced dedifferentiation (Liu et al., 2015). Notable is the fact that not only Ptgs2 but also Ptgs1 is significantly induced alongside blastema formation in reptiles (Xu et al., 2019). Thus, OSKM-mediated dedifferentiation would be accounted for by the anomalous induction of Ptgs1 by partial reprogramming, possibly via epigenetic alterations. As a follow-up study, it would be of interest to determine the epigenetic profile of Ptgs1 to account for the OSKM-induced Ptgs1 expression in light of the drastic epigenetic alterations induced by partial reprogramming (Browder *et al*., 2022b).

## Methods

### Mice

*Col1A1*^*TetO-OSKM*^;*ROSA26*^*rtTA*^ mice were obtained from Jackson Laboratory (#011004). *ROSA26*^*rtTA*^ mice were generated from *Col1A1*^*TetO-OSKM*^;*ROSA26*^*rtTA*^ mice. Age-matched mice between 2-4 months were utilized for all experiments. For OSKM induction, mice were treated with 0.15mg/ml doxycycline in drinking water containing 5% sucrose for 4 days. For BrdU pulse-chase experiment, BrdU (150mg/kg) was intraperitoneally injected 3 hours before sacrifice. For injury experiments, mice were treated with 10Gy X-ray radiation 2 days after doxycycline treatment and sacrificed at indicated time points. For the inhibition of prostaglandin production while OSKM induction mice were treated with 0.15mg/ml doxycycline in drinking water containing 5% sucrose for 4days and addition of 0.6mg/ml dexibuprofen in the last 2days. For Cox-1,2 inhibition, SC-560 (15mg/kg/day) and Celecoxib (18.75mg/kg/day) were intraperitoneally injected respectively for the last two days of doxycycline administration. These animal experiments were conducted under the permission of Seoul National University Institutional Animal Care and Use Committee (Permission number: SNU-210326-5-4, SNU-221017-2).

### *In vitro* culture of intestinal organoid

For *in vitro* culture of adult organoids, mouse small intestine was washed with PBS and cut longitudinally. Villi were scraped away with a microscope slide and the intestine was washed three times with cold PBS. The intestine was then incubated in gentle cell dissociation reagent (STEMCELL technologies, #07174) for 15 min. The isolated crypts were embedded in Matrigel (Corning, 354234) and seeded on 48-well plate. After polymerization, intesticult™ (STEMCELL technologies, #06005) containing 50μg/ml gentamicin was added and refreshed every 2 days. For passaging, the organoids were suspended in cold DMEM/F12 (Gibco) and were embedded in fresh Matrigel and seeded on plate followed by addition of culture medium. For OSKM induction, organoids were treated with 0.1 or 0.5μg/ml doxycycline for 3 days. For BrdU pulse-chase experiment, organoids were treated with 10μM BrdU for 5 hours. For injury experiments, organoids were treated with 6Gy or 10Gy X-ray radiation. For the inhibition of prostaglandin production while OSKM induction, organoids were treated with 0.5μg/ml doxycycline alone for 1 day and with 3μM Dexibuprofen addition for 2 days. For the inhibition of Cox-1 and Cox-2, organoids were treated with 3μM Celecoxib and 0.675μM SC-560, respectively. For *in vitro* culture of fetal organoids, fetal small intestines were cut into small pieces, and then dissociated with gentle cell dissociation reagent (STEMCELL technologies) for 15 min. The isolated epithelial units were embedded in Matrigel and maintained in conditions identical to those used for adult intestinal organoids.

### Immunohistochemistry and Immunofluorescence

For intestine sections, Swiss rolls of mouse intestines were fixed in 4% PFA at 4°C overnight, immersed in 30% sucrose at 4°C overnight, and then cryopreserved in OCT compound (Sakura Finetek). For organoid sections, intestinal organoids were fixed in 4% PFA at room temperature for 15 min, immersed in 30% sucrose at 4°C overnight, and then cryopreserved in OCT compound. For immunohistochemistry, 6-μm sections were treated with 3% H_2_O_2_ in methanol for blocking endogenous peroxidase activity. For antigen retrieval. slides were kept in Tris-EDTA buffer (pH 9.0) at 95°C for 20 min. Nonspecific binding was prevented by incubation with 10% FBS in PBS for 1 hour. slides were incubated with a primary antibody at 4°C overnight or at room temperature for 1 hour. Then, the slides were incubated with a horseradish peroxidase-conjugated secondary antibody at room temperature for 2 hours, followed by 5 to 10 min incubation with DAB substrate (Vector lab., SK-4100). Hematoxylin solution (Sigma Aldrich, GHS3) was used for nuclear staining and then washed with tap water. slides were covered with slide glass using MOWIOL solution. Samples were visualized with a microscope (Leica DM500). For immunofluorescence, sections were incubated with a primary antibody in 4% BSA in PBS at 4°C overnight or at room temperature for 1 hour. Then, the slides were incubated with Alexa Fluor secondary antibodies conjugated to 488 or 594 fluorophores (Invitrogen) at room temperature for 2 hours. DAPI (Thermo Fisher) was used for nuclear staining and then washed with PBS-T. slides were covered with slide glass using MOWIOL solution. Fluorescence microscopy (Olympus) was used for imaging samples. Following primary antibodies were used for IHC and IF: mouse anti-2A peptide (Millipore, MABS2005, 1:200), rabbit anti-Active YAP1 (Abcam, ab205270, 1:1000), rabbit anti-Avil (Abcam, ab72210, 1:500), rabbit anti-BrdU (Novus Biologicals, NBP2-14890, 1:200), CD44 (BioLegend, 103015, 1:500), rabbit anti-Ki67 (Abcam, ab16667, 1:500), rabbit anti-Lysozyme (Abcam, ab108508, 1:200), mouse anti-Oct4 (BD Biosiences, 611203, 1:500), rabbit anti-Olfm4 (CST, 39141, 1:400), rabbit anti-Phosphohistone H3 (CST, 53348, 1:1000), rat anti-Sca1 (BioLegend, 122501, 1:500), rabbit anti-Sox2 (Millipore, AB5603, 1:400), goat anti-Trop2 (R&D Systems, AF1122, 1:100), mouse anti-Cox1 (Santa cruz, sc-19998, 1:100), mouse anti-Cox2 (Santa cruz, sc-19999, 1:100), and rabbit anti-Dclk1(Abcam, ab31704, 1:1000).

### Histology and AB-PAS Staining

For histological analysis, sections were stained with hematoxylin and eosin, dehydrated, and then covered with slide glass using Canada balsam (Sigma Aldrich). For staining of goblet cells, sections were stained using Alcian blue(AB) and periodic acid-Schiff (PAS). Samples were visualized with a microscope (Leica DM500).

### In situ hybridization

Intestine were fixed overnight in 4 % paraformaldehyde in phosphate-buffered saline (PBS). For whole mount in situ hybridization, the intestine was treated with 20 μg/ml proteinase K (AM2546, Thermo Fisher Scientific, United States) for 10 min at room temperature. Antisense RNA probes were labeled with digoxigenin (Roche, Switzerland). Digoxigenin (DIG)-labeled RNA probes were pre-warmed at 85°C and hybridized to the intestine specimen overnight at 69°C. After whole mount in situ hybridization, the specimens were cryo-sectioned at a thickness of 12 μm. At least 10 specimens were examined for each condition. The primer sequences of the Clu and Msln are as follows: Clu-Forward: 5’-GAG ATT CAG AAC GCC GTC CA-3’ Clu-Reverse: 5’-CTC TTG TGT GGG AAG CCG AT-3’ Msln-Forward: 5’-GTG CCC ACT TCT TCT CCC TC-3’ Msln-Reverse: 5’-GGT GCC ATC TAC ACA AGC CT-3’ For Clu in situ hybridization in Figure 6, RNAscope™ 2.5 HD Assay – RED (ACD, 322360), RNAscope® Probe-Mm-Clu (ACD, 427891), and RNAscope® H202 & Protease Plus Reagents (ACD, 322330) were used with mouse intestinal sections prepared as for IFC and IHC, following provided protocol (Document Number 322360-USM).

### RNA isolation and quantitative RT-PCR

Easy-BLUE™ RNA isolation kit (iNtRON Biotechnology, #17061) is used for total RNA extraction. PrimeScript™ RT reagent kit (TaKaRa, RR036A) is used to generate cDNA from RNA extracted. Quantitative real-time PCR analysis was performed with Light Cycler-480^®^II (Loche) using TB-Green (Takara, RR420) following the supplier’s instructions.

### Immunoblotting

Cell lysates were extracted with RIPA buffer supplemented with 1% protease inhibitor cocktail and 0.1% sodium orthovanadate. After 1 hour incubation on ice, total protein was extracted after centrifugation. The concentration of total protein was quantified by BCA protein assay kit (Thermo Scientific). 10μg of total protein was separated on 10% SDS/PAGE. Separated protein in the gel was transferred to PVDF membrane. Membrane with protein was blocked with 10% skim milk in Tris-buffered saline containing 0.1% Tween-20 (TBS-T) for 1 hour and then washed three times by TBS-T. The membrane was incubated with primary antibody in TBS-T at 4°C overnight. Following primary antibodies were used: mouse anti-Oct4 (BD Biosiences, 611203, 1:1000), and mouse anti-a-tubulin (Santa cruz, sc-8035, 1:1000). Incubated membrane was washed three times with TBS-T. The membrane was incubated at room temperature with HRP-conjugated secondary antibody in TBS-T for 1 hour, followed by wash with TBS-T three times. Immunoreactivity was detected by Chemi-Doc using WEST-Queen™ kit (iNtRON Biotechnology, #16026).

### Flow Cytometry

Intestinal organoids were incubated in TrypLE (Invitrogen) for 15 min at 37°C for single cell dissociation and then washed with PBS three times. For Sca1 staining, the dissociated cells were incubated in PE/Cy7-conjugated anti-Sca1 antibody (BD Biosciences, 558162, 1:100) in 4% BSA in PBS for 20 min. Then, the cells were washed three times and analysed by Flow cytometer (BD LSRFortessa X-20). The data were analysed with FlowJo software.

### Caspase-3 activity assay

Caspase□3 activity was detected using colorimetric assay kits (Abcam, ab39401). The kits were used according to the manufacturer’s protocols. Briefly, 24 hours after IR, organoids were lysed in the supplied lysis buffer for 10 min at 4°C. Supernatants were collected and incubated with the supplied reaction buffer containing dithiothreitol and DEVD□p-NA substrate at 37°C for 2 hours. The reactions were measured by changes in absorbance at 400 nm using a microplate reader.

### Mouse intestinal epithelial and mesenchymal cells isolation

Mouse intestine was dissected and washed with PBS and cut longitudinally. Intestine was incubated in PBS containing 2mM EDTA for 30 min, followed by vigorous shaking and washed with PBS. For mesenchymal cells isolation, epithelial cells were removed as mentioned above, and incubated in DMEM containing 10% FBS, Collagenase XI (300 units/ml), Dispase I (2 units/ml) and DNase II (50 units/ml) for 1 h, at 37□°C. Isolated cells were washed with 2% sorbitol containing Red Blood Cell Lysis Solution.

### Prostaglandin E2 assay

Epithelial and mesenchymal cells were isolated as mentioned above with every reagent containing indomethacin (6μg/ml). Isolated cells were sonicated for 3 min using cycles of 2 seconds, followed by centrifugation for 5 min, 13000 rpm. Using PGE2 ELISA kit (R&D Systems, # KGE004B), Prostaglandin E2 level was measured and standardized with protein level using BCA Protein Assay Kit (Thermo Fisher, #23227).

### Single-cell RNA-seq library preparation

To prepare single cells, whole epithelial segments were isolated with 2mM EDTA for 30 min, washed with PBS, and then dissociated to single cells with TrypLE (Invitrogen) at 37°C for 30 min, with mixing every 10 min. Dissociated cells were filtered through a 40-μm cell strainer on ice and washed three times with cold DMEM/F12. Single-cell suspensions with 90% cell viability were processed on the 10x Chromium Controller using Chromium Next GEM Single Cell 3’ Reagent Kits v3.1 (10x Genomics) according to the manufacturer’s instructions. Cells were partitioned into nanoliter-scale Gel Beads-in-Emulsion (GEMs) with target recovery of 10,000 cells. The single-cell 3 prime mRNA seq library was generated by reverse transcription, cDNA amplification, fragmentation, and ligation with adaptors followed by sample index PCR. Resulting libraries were quality checked by Bioanalyzer and sequenced on an Illumina NovaSeq 6000 (index = 8 bases, read 1 = 26 bases, and read 2 =□91 bases).

### scRNA-seq data preprocessing

We processed raw FASTQ files for scRNA-seq using the CellRanger software suite (v5.0.1) (Zheng et al., 2017). Reads were mapped to the mouse reference genome (GRCm38) with the Ensembl GRCm38.102 GTF file. We also eliminated technical artifacts using the remove-background function of the CellBender (v0.2.0) python package (Fleming et al., 2019). We excluded low quality cells with < 2.0 log10-scaled counts of unique molecular identifiers (UMIs) and < 50% of UMIs assigned to mitochondrial genes using the calculateQCMetric function of the scater (v1.22.0) R package (McCarthy et al., 2017). To remove cell-specific biases, cells were grouped using the quickCluster function of the scran (v1.22.0) R package(Lun et al., 2016) with default options and cell-specific size factors were calculated using the computeSumFactors function of the same package with default options. Raw counts of each cell were divided by cell-specific size factor and log2-transformed with the pseudocount of 1. To define highly variable genes (HVGs), we modeled the variance of the log-expression profiles of each gene and decomposed it into technical and biological components based on a fitted mean-variance trend using the modelGeneVar function of the scran R package. Genes with < 0.05 FDR were defined as HVGs. For downstream analysis, we calculated the top 20 principal components (PCs) calculated from the normalized count matrix of HVGs. To remove technical effects between different samples, calculated principal components were corrected using the RunHarmony function of the harmony (v0.1.0) R package (Korsunsky et al., 2019). A Shared Nearest Neighbor (SNN) graph was constructed using the FindNeighbors function of the Seurat (v4.0.5) R package (Butler et al., 2018) with the top 20 PCs and cells were clustered using the FindClusters function with the default options based on a SNN graph. Cells were visualized on the two-dimensional UMAP plot using the RunUMAP function of the same package as with the first 20 PCs. Each cluster was defined as one of the major cell types based on the expression of canonical cell-type marker genes while clusters identified as non-epithelial cells – including Ptprc+ immune cells – were excluded for downstream analysis. The remaining cells were re-grouped and visualized on the 20 PCs using the same method described above. For projection, reference data (revSC and aVEC-containing scRNA-seq data, GSE123516, GSE169718, respectively) were preprocessed using the same methods as above. We annotated the major cell types mentioned in the paper based on the expression of canonical cell type-specific marker genes after excluding non-epithelial cells including Ptprc+ immune cells. For our scRNA-seq and reference scRNA-seq data, differentially expressed genes (DEGs) of each cell type were computed using the FindMarkers function of Seurat R package.

### scRNA-seq data analysis

To infer the stemness of each cell, we estimated differentiation potency using the SCENT (v0.3.3) R package (Chen and Teschendorff, 2019). A raw count matrix log-transformed with a pseudocount of 1.1 to avoid 0 values after log-transformation and used as an input matrix. Normalized data was integrated with the PPI network using the DoIntegPPI function and signaling entropy rate (SR) was computed using the CompSRana function with default options. Additionally, we inferred differentiation potency using the CytoTRACE function of the CytoTRACE (v1.0.2) R package (Gulati et al., 2020) with default options. For 1,275 fetal signature genes (Yui et al., 2018b) and 398 yap signature genes (Barry et al., 2013), gene signature score per cell was calculated using the AddModuleScore function of Seurat R package. To project reference data (revSC and aVEC-containing scRNA-seq data) onto our data, for each reference data, we obtained k-Nearest Neighbors (k-NNs) from our data based on Pearson correlation coefficients of normalized expression data of HVGs between the reference data and our data using the knn.index.dist function of the KernelKnn (v1.1.4) R package (Mouselimis, 2021). To visualize the projection results, for each reference data, we averaged two-dimensional coordinates of 10-NNs on the UMAP plot of our data. To infer the orientation of cellular differentiation, we performed RNA velocity analysis using the scVelo (v0.2.4) python package (Bergen et al., 2020). For each condition, we generated the spliced and unspliced expression profiles using the run function of the velocyto (v0.17.17) python package (La Manno et al., 2018) with the mouse reference masking GTF file (GRCm38) and the Ensembl GRCm38.102 GTF file. The raw profiles were natural log-transformed after excluding low expressed genes using the pp.filter_and_normalize function with default parameters except for min_shared_counts=30. Moments for velocity estimation were computed using the pp.moments function with the default parameters except for n_neighbors=5. RNA velocities were estimated using the tl.velocity function with the option of mode=dynamical. The velocity graphs were computed based on cosine similarities using the tl.velocity_graph function with default options. We visualized RNA velocity results on the two-dimensional t-SNE plot using the Palantir (v1.0.0) python package (Setty et al., 2019). For the Palantir t-SNE plot, we computed diffusion components (DCs) using the run_diffusion_maps function with the first 200 PCs. A k-Nearest Neighbor (kNN) graph (k=30) was constructed from the first 20 DCs. The coordinated for t-SNE plot were computed using the run_tsne function with the options of perplexity=300. To quantify the connectivity between the 19 cell clusters, we generated partition-based graph abstraction (PAGA) graph of each condition using the tl.paga function with default options. The PAGA graphs with velocity-directed edges were plotted using only directed arrows.

### Bulk RNA-seq library preparation

Total RNA was isolated from intestinal organoids using Easy-BLUE™ RNA isolation kit (iNtRON Biotechnology, #17061). One 1 µg of total RNA was processed for preparing mRNA sequencing library using MGIEasy RNA Directional Library Prep Kit (MGI) according to manufacturer’s instruction. The first step involves purifying the poly-A containing mRNA molecules using poly-T oligo attached magnetic beads. Following purification, the mRNA is fragmented into small pieces using divalent cations under elevated temperature. The cleaved RNA fragments are copied into first strand cDNA using reverse transcriptase and random primers. Strand specificity is achieved in the RT directional buffer, followed by second strand cDNA synthesis. These cDNA fragments then have the addition of a single ‘A’ base and subsequent ligation of the adapter. The products are then purified and enriched with PCR to create the final cDNA library. The double stranded library is quantified using QauntiFluor ONE dsDNA System (Promega). The library is circularized at 37 °C for 30 min, and then digested at 37 °C for 30 min, followed by cleanup of circularization product. To make DNA nanoball (DNB), the library is incubated at 30 °C for 25 min using DNB enzyme. Finally, Library was quantified by QauntiFluor ssDNA System (Promega). Sequencing of the prepared DNB was conducted on the MGIseq system (MGI) with 100 bp paired-end reads.

### Bulk RNA-seq processing and analysis

Low-quality bases and adapter sequences bases were trimmed using TrimGalore (https://www.bioinformatics.babraham.ac.uk/). The trimmed reads were aligned to the mouse genome assembly GRCm39 using STAR (v2.7.3a). Read counts were normalized by trimmed mean of M-values and log_2_ fold changes (log2FC) of genes between conditions were obtained by using the DESeq function of DESeq2 (v1.34.0) R package. Genes with log2FC > 0.5 and adjusted P-values < 0.05 were identified as differentially expressed genes (DEGs). Functional enrichment analysis of DEGs for “WikiPathway_2021_Human” database was performed using the enrichR (v3.1) R package. To compare *Ptgs1* and *Ptgs2* mRNA expressions without any technical biases, three expected read count matrices were combined and normalized using the same methods as described above.

### Quantification and statistical analysis

The quantitative data are expressed as the mean values ± standard deviation (SD). Student’s unpaired t-tests was performed to analyze the statistical significance of each response variable using the PRISM. p values less than 0.05 were considered statistically significant, p < 0.05(*), p <0.01(**), p <0.001(***), and p <0.0001(****).

## Data Availability

All unique/stable reagents generated in this study will be freely available from the lead contact to academic researchers with a completed Materials Transfer Agreement.

The bulk RNA-seq and scRNA-seq data are available from Sequence Read Archive under accession numbers: PRJNA854284 and PRJNA853815, respectively.

## Acknowledgments

This work was supported by a grant from the National Research Foundation of Korea (NRF-2020R1A2C2005914, NRF□2019M3A9D5A01102794, 2021M3H9A1030158, NRF-2016R1A5A2008630 and NRF-2020M3A9E4037904), by Korean Fund for Regenerative Medicine funded by Ministry of Science and ICT, and Ministry of Health and Welfare (Grant number RS-2022-00070316).

## Author contributions

H-J. C. and B-K. K conceived the overall study design, led the experiments, and wrote the manuscript. J.K. and S-Y.L, mainly conducted the experiments, data analysis, and wrote the first draft. C.P.H., and S.K., performed scRNA-seq and bulk RNA-seq data analysis, and wrote the specific part of the manuscript. J-K. K., led the analysis of scRNA-seq data and wrote the manuscript. B-K. J., J-Y. O., and K-T. K., conducted the repeat experiments and validated the results. E-J. K., performed the bulk RNAseq data analysis. A. A. A., E-J. K., and H-S. J. conducted ISH assay.

