## Supplemental figures and legends for "Partial *in vivo* reprogramming enables injury-free intestinal regeneration via autonomous Ptgs1 induction"

**Running title: Ptgs1 for injury-free intestinal regeneration**

Jumee Kim, Somi Kim, Seung-Yeon Lee, Beom-Ki Jo, Ji-Young Oh, Eun-Ji Kwon, Keun-Tae Kim, Anish Ashok Adpaikar, Eun-Jung Kim, Han-Sung Jung, Chang Pyo Hong, Jong Kyoung Kim, Bon-Kyoung Koo, Hyuk-Jin Cha

This PDF file includes:

Experimental Section

Supplemental Figure legends

Supplementary Figures 1-6

**Figure S1. Identification of dedifferentiated intestinal epithelial cells by partial reprogramming**

(A) Relative mRNA expression of OSKM in intestinal epithelial cells of iOSKM mice 4 days after Dox treatment. Data represent the mean with SD (n = 6; 3 mice x 2 technical replicates) (B) Necropsy of iOSKM mice 4 days after Dox treatment. (C) Body weight of iOSKM mice during Dox treatment. Data represent the mean with SD (n = 3 mice) (D) IF of Ki67 in the intestine (left) and quantification of Ki67-positive cells per crypt (-Dox, n = 10; +Dox, n = 13) (right). (E) Violin plots showing log10-scaled value of unique molecular identifiers (UMIs) (left) and log10-scaled value of the number of features in –Dox and +Dox conditions. (F) UMAP plot of scRNA-seq from OSKM-induced mouse intestinal epithelium indicated with 18 distinct clusters. (G) Dot plot for expression of canonical marker genes per each cluster. (H) Bar plot showing cell type proportion in each condition. (I) Box plot indicating signaling entropy rate (SR) inferred by SCENT per each condition in different cell types. (J) Violin plots showing expression of differentiated cell-type marker genes between conditions in different cell types. (K) IF of Trop2 in the intestine of iOSKM mice. DAPI for nuclear staining. Data represent the mean with SD. Student's t-test: $p < 0.05$(*), $p < 0.01$(**), $p < 0.001$(***), $p < 0.0001$(****), ns, not significant. Scale bar = 50μm.

**Figure S2. Concurrent fetal gene transition and YAP activation by partial reprogramming in intestinal organoids**

(A) IF of Oct4 and 2A peptide in iOSKM intestinal organoids. DAPI for nuclear staining. (B) Relative mRNA expressions of OSKM in intestinal organoids. (C) Relative mRNA expressions of *Nanog*, pluripotency marker, in intestinal organoids. (D) Microscopic images of Dox-treated intestinal organoids (left) and quantification of spheroid/budding organoid ratio (right) (n =4).

(E) Microscopic images of intestinal organoids with Dox treatment for 2days, with Dox treatment for 7 days, and with Dox treatment for 2 days followed by Dox removal for 5 days. (F) Microscopic images and IF of fetal genes, Trop2 and Sca1, in intestinal organoids from maternal and E17.5 fetal mice. (G) Flow cytometry of Sca1 in Dox-treated intestinal organoids. (H) Relative mRNA expressions of fetal genes, *Ly6a* (encoding Sca1), *Anxa1*, and *Tacstd2* (encoding Trop2), in iOSKM intestinal organoids. (I) Relative mRNA expressions of Hippo pathway-associated genes, *YAP*, *Ccn2* (encoding Ctgf), and *Tead4*, in intestinal organoids. (J) Heatmap showing relative OSKM expression in intestinal organoids.

**Figure S3. Characterization of OSKM-induced revSC-like and aVEC-like cells**

(A) UMAP plots showing conditions (left) and cell types (middle) of revSC-containing scRNA-seq data and projection results of revSC-containing scRNA-seq data onto our scRNA-seq data (right). (B) UMAP plots showing conditions (left) and cell types (middle) of aVEC-containing scRNA-seq data and projection results of aVEC-containing scRNA-seq data onto our scRNA-seq data (right). (C) Relative expression of revSC, aVEC, EC and Fetal in each cell type, (D-F) Violin plots showing expression for revSC (D), aVEC (E) marker genes, and DC2-specific expressed genes (F). (G) Venn diagrams showing overlapping cell type-specific marker genes between DC1, DC2 and EC. Colors indicating reported revSC-specific (red), revSC/aVEC common (purple) and aVEC-specific (blue) marker genes. (H) Violin plots showing expression for villus-top (left) and villus-bottom (right) EC marker genes. (I) UMAP plots showing gene expression of *Ly6a* (encoding Sca1) and *Avil*. (J) t-SNE plots showing RNA velocity inferred by scVelo in –Dox and +Dox conditions.

**Figure S4. Promoted intestinal regeneration after IR damage by partial reprogramming**

(A) Western blotting for Oct4 in intestinal epithelial cells of iOSKM mice on Dox treatment for indicated day(s). $\alpha$-tubulin for loading control. (B, C) Intestine of iOSKM mice after 10Gy IR. Dox was treated 2 days before IR for 4 days. H&E histology at 4dpi (B) and IF of Trop2 at

2dpi and 4dpi (C). (D) Caspase-3 activity using intestinal epithelial cells 24 hours after IR. ns, not significant. (E) Microscopic images of intestinal organoids with Dox and IR treatment. The morphology was analyzed at day after irradiation as indicated.

**Figure S5. Inhibition of fetal gene transition and YAP activation by NSAID in intestinal organoids**

(A) Bar plots showing enrichR combined scores of pathways commonly upregulated in indicated condition (B) Microscopic images of Control (left, left) Dox-(left, middle), Dox and NSAID-(left, right) treated intestinal organoids and quantification of spheroid/budding organoid ratio (right) (n =4). (C) IF of Sca1, in mouse intestinal organoids. (D) mRNA expressions of fetal genes, *Ly6a* (encoding Sca1), iOSKM intestinal organoids. (E) IF of active YAP, in mouse intestinal organoids. DAPI for nuclear staining

**Figure S6. Increase in Cox1 expression and prime role in intestinal regeneration**

(A) IF of Cox1 and Dclk1(a marker or tuft cells) in the intestine of iOSKM mice. (B) Normalized mRNA expressions of *Ptgs1* and *Ptgs2* (encoding Cox1, Cox2 respectively) in iOSKM intestinal organoids (n = 8; 4 x 2 technical replicates). (C) IF of Cox1 and Cox2 in iOSKM intestinal organoids. DAPI for nuclear staining. (D) IF of Sca1, Avil and Cox1 in iOSKM intestinal organoids. (E) IF of Sca1 in Control, Dox treated, Dox and iCox1, Dox and iCox2 treated (from left to right) iOSKM intestinal organoids.

**Movie S1.** Growth of intestinal organoids in both conditions, control (A) and Dox treatment (B), for 3 days

#### Figure S1.

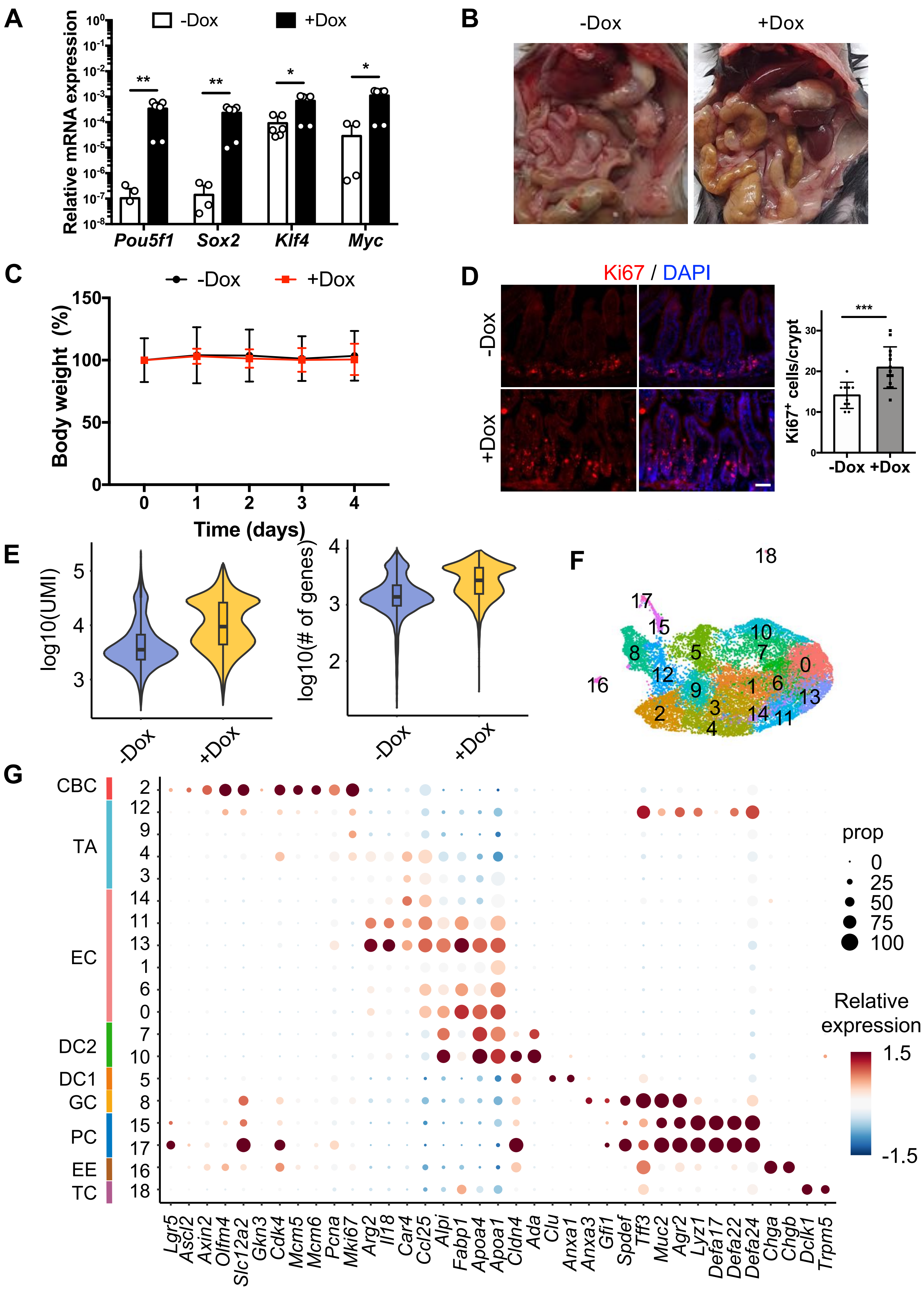

A
-Dox
+Dox
Relative mRNA expression
Pou5f1
Sox2
Klf4
Myc
B
-Dox
+Dox
C
-Dox
+Dox
Body weight (%)
Time (days)
D
Ki67 / DAPI
-Dox
+Dox
Ki67+ cells/crypt
E
log10(UMI)
log10(# of genes)
-Dox
+Dox
F
G
CBC
TA
EC
DC2
DC1
GC
PC
EE
TC
prop
Relative expression
Lgr5
Ascl2
Axin2
Olfm4
Slc12a2
Gkn3
Cdk4
Mcm5
Mcm6
Pcna
Mki67
Arg2
Il18
Car4
Ccl25
Alpi
Fabp1
Apoa4
Apoa1
Cldn4
Ada
Clu
Anxa1
Anxa3
Gfi1
Spdef
Tff3
Muc2
Agr2
Lyz1
Defa17
Defa22
Defa24
Chga
Chgb
Dclk1
Trpm5

#### Figure S1.

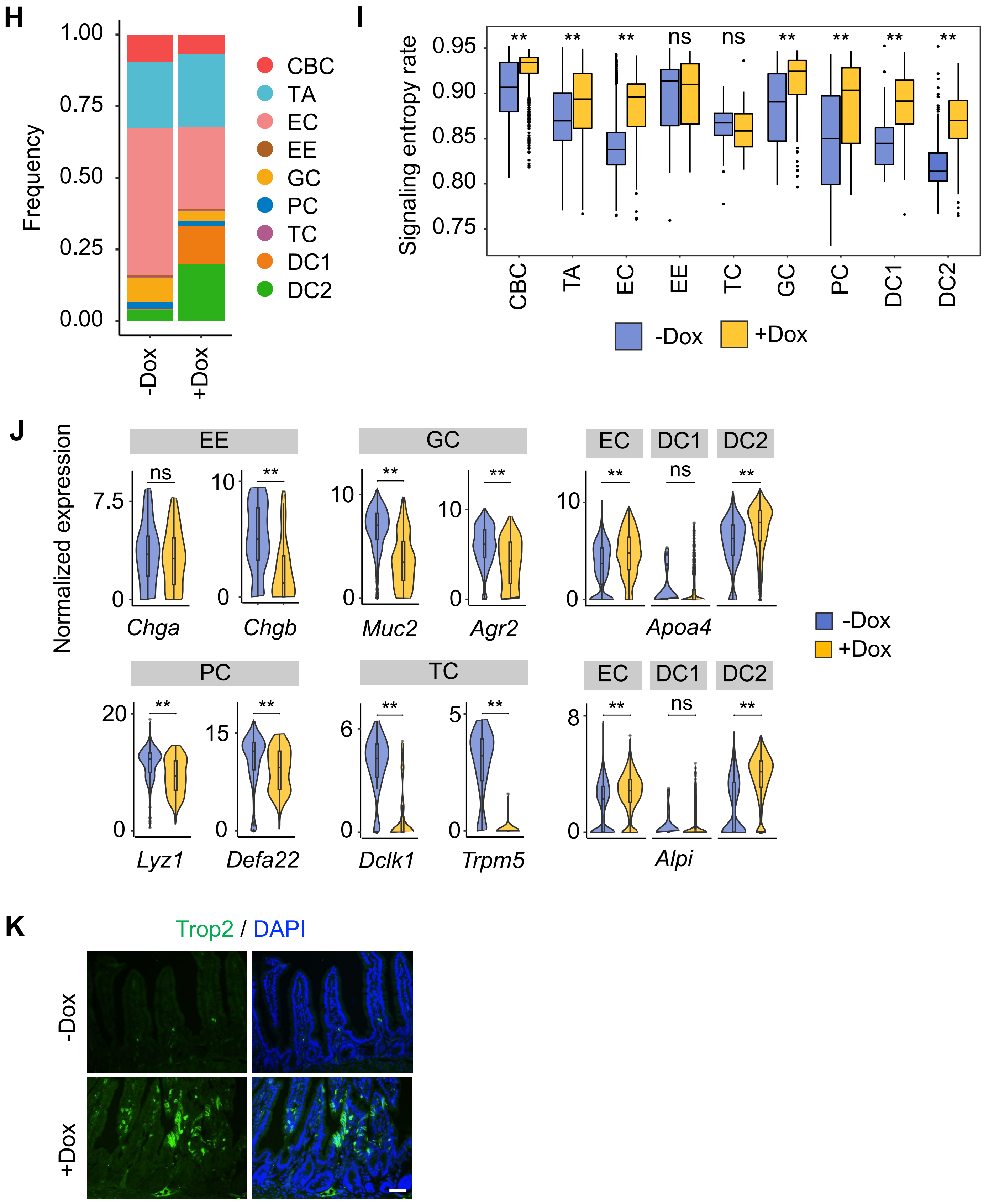

H
1.00
0.75
0.50
0.25
0.00
Frequency
-Dox
+Dox
CBC
TA
EC
EE
GC
PC
TC
DC1
DC2
I
Signaling entropy rate
0.95
0.90
0.85
0.80
0.75
** ** ** ns ns ** ** ** **
CBC TA EC EE TC GC PC DC1 DC2
-Dox
+Dox
J
Normalized expression
EE
GC
EC DC1 DC2
ns ** ** ** ** ns **
Chga Chgb Muc2 Agr2 Apoa4
-Dox
+Dox
PC
TC
EC DC1 DC2
** ** ** ** ** ns **
Lyz1 Defa22 Dclk1 Trpm5 Alpi
K
Trop2 / DAPI
-Dox
+Dox

**Figure S2.**

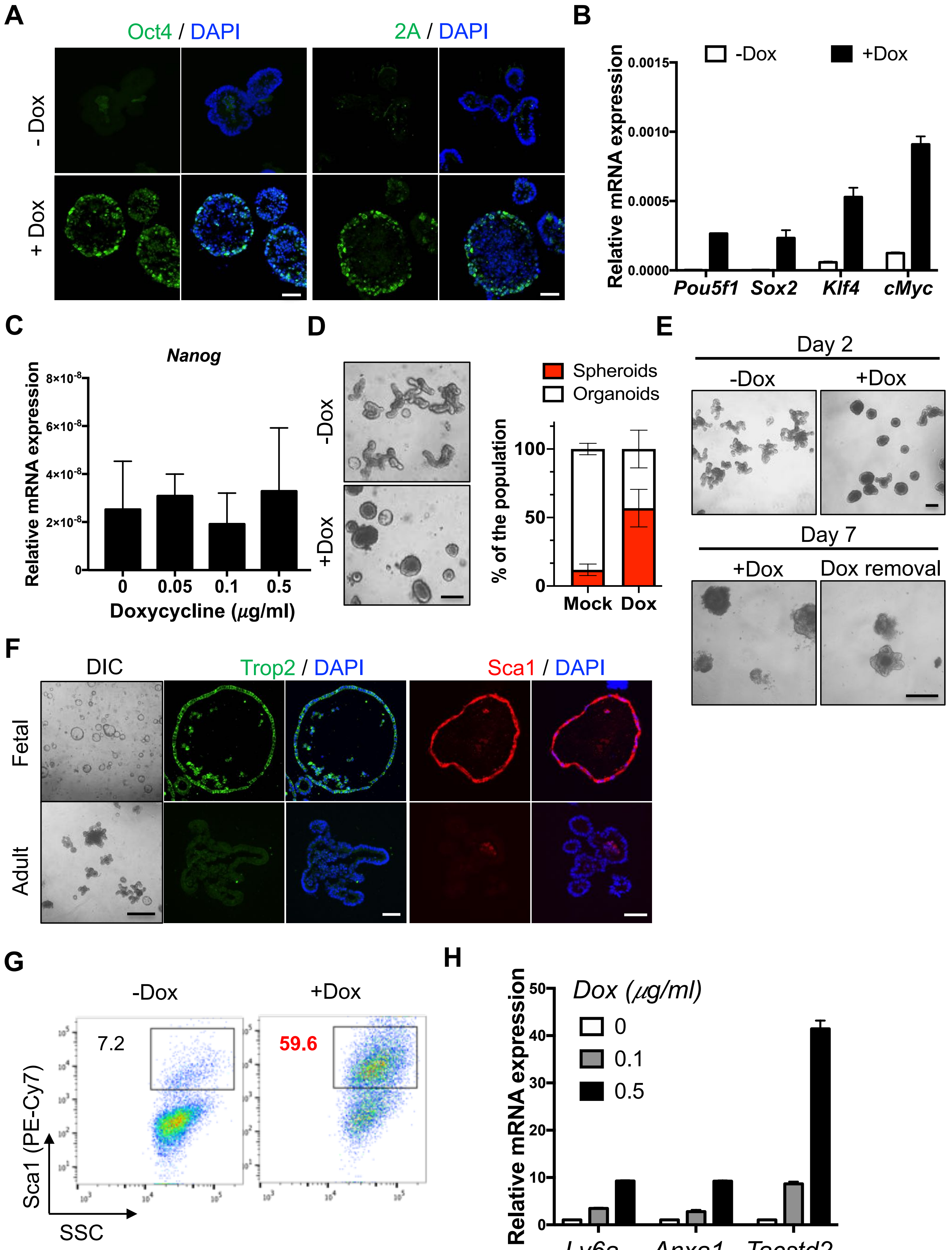
A
Oct4 / DAPI
2A / DAPI
- Dox
+ Dox
B
Relative mRNA expression
-Dox
+Dox
0.0015
0.0010
0.0005
0.0000
Pou5f1
Sox2
Klf4
cMyc
C
Nanog
Relative mRNA expression
8×10-8
6×10-8
4×10-8
2×10-8
0
0
0.05
0.1
0.5
Doxycycline (μg/ml)
D
-Dox
+Dox
Spheroids
Organoids
% of the population
100
50
0
Mock
Dox
E
Day 2
-Dox
+Dox
Day 7
+Dox
Dox removal
F
DIC
Trop2 / DAPI
Sca1 / DAPI
Fetal
Adult
G
-Dox
+Dox
7.2
59.6
Sca1 (PE-Cy7)
SSC
H
Relative mRNA expression
Dox (μg/ml)
0
0.1
0.5
50
40
30
20
10
0
Ly6a
Anxa1
Tacstd2

**Figure S2.**

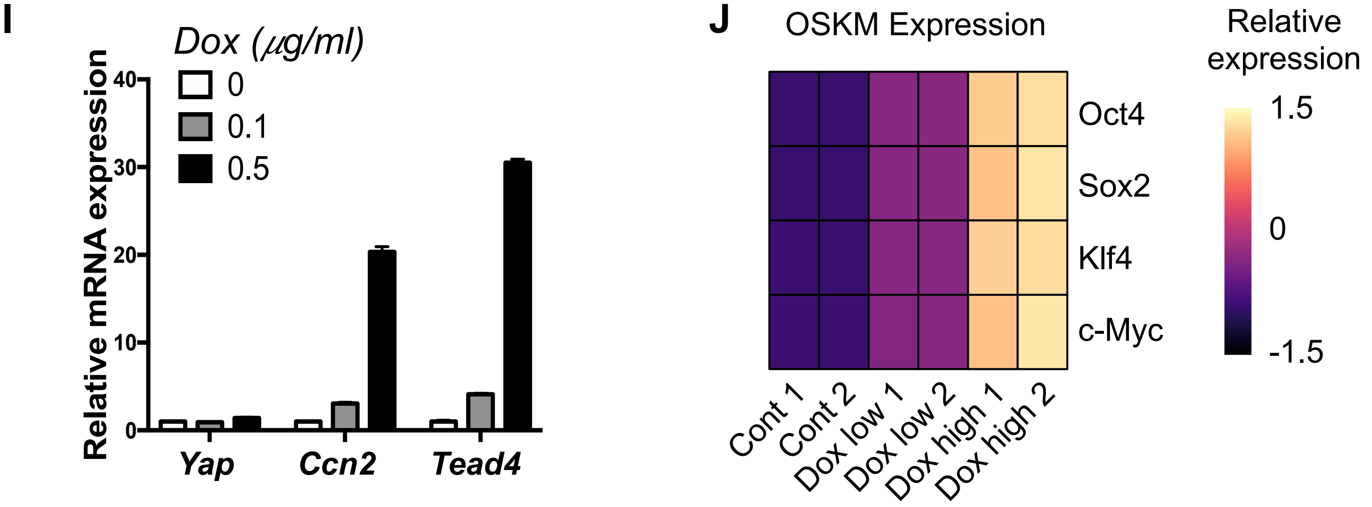

I
Dox (μg/ml)
0
0.1
0.5
Relative mRNA expression
40
30
20
10
0
Yap
Ccn2
Tead4
J
OSKM Expression
Relative
expression
1.5
0
-1.5
Oct4
Sox2
Klf4
c-Myc
Cont 1
Cont 2
Dox low 1
Dox low 2
Dox high 1
Dox high 2

Figure S3.

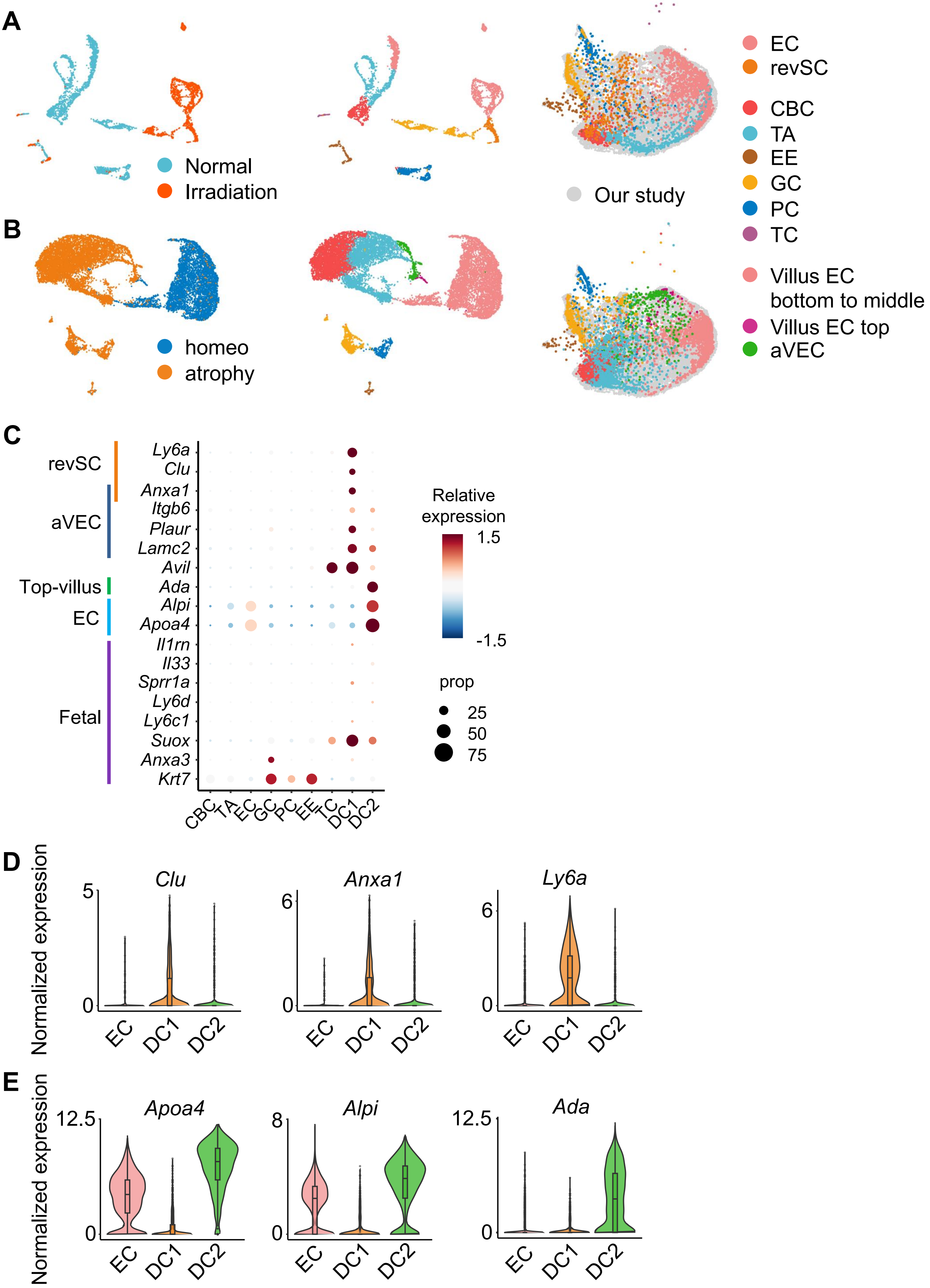
A
Normal
Irradiation
Our study
EC
revSC
CBC
TA
EE
GC
PC
TC
B
homeo
atrophy
Villus EC
bottom to middle
Villus EC top
aVEC
C
revSC
aVEC
Top-villus
EC
Fetal
Ly6a
Clu
Anxa1
Itgb6
Plaur
Lamc2
Avil
Ada
Alpi
Apoa4
Il1rn
Il33
Sprr1a
Ly6d
Ly6c1
Suox
Anxa3
Krt7
CBC
TA
EC
GC
PC
EE
TC
DC1
DC2
Relative
expression
1.5
-1.5
prop
25
50
75
D
Normalized expression
Clu
Anxa1
Ly6a
EC
DC1
DC2
E
Normalized expression
Apoa4
Alpi
Ada
12.5
8

**Figure S3.**

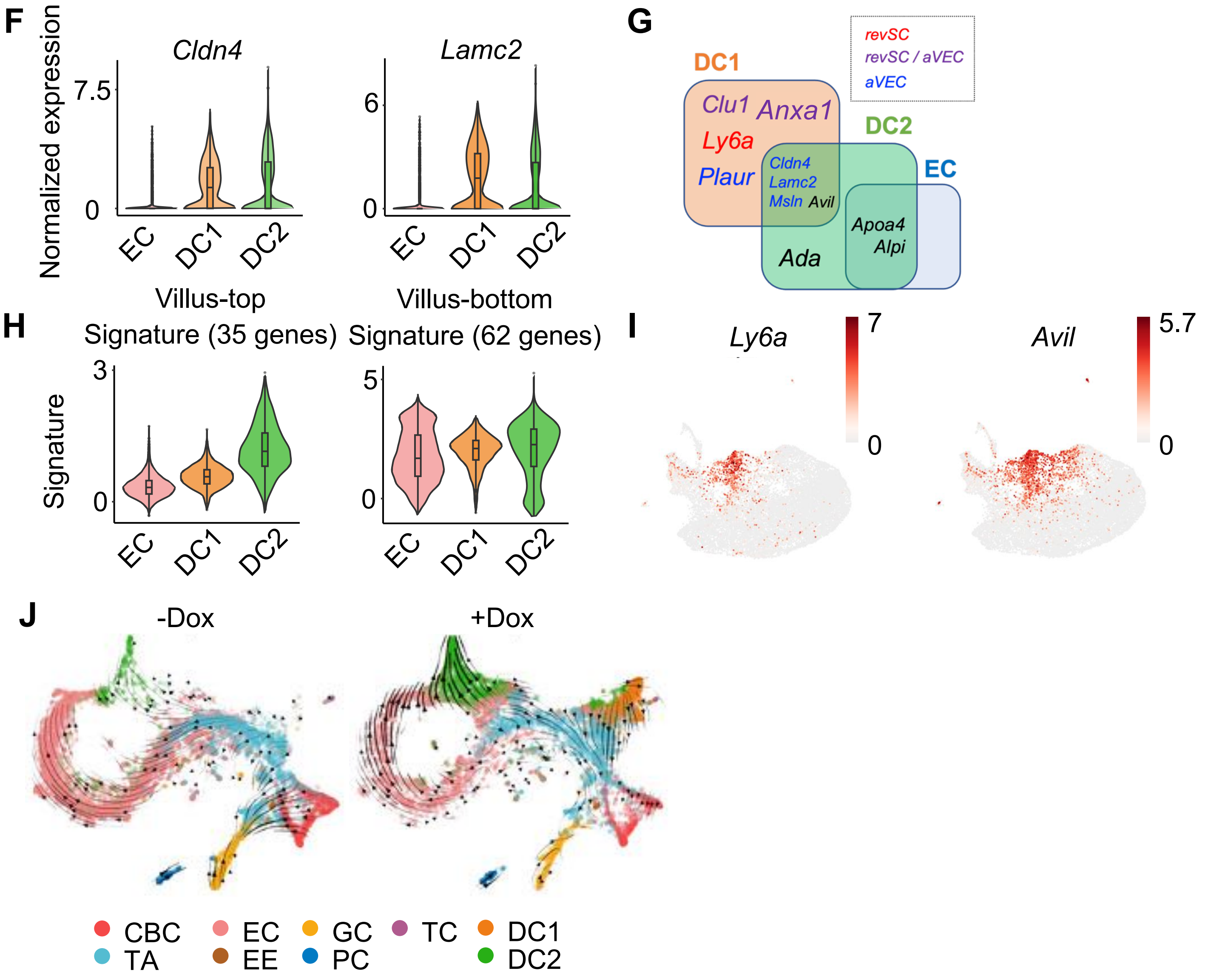

F
Cldn4
Lamc2
Normalized expression
7.5
0
6
0
EC
DC1
DC2
EC
DC1
DC2
G
revSC
revSC / aVEC
aVEC
DC1
Clu1
Anxa1
Ly6a
Plaur
DC2
Cldn4
Lamc2
Msln
Avil
EC
Apoa4
Alpi
Ada
H
Villus-top
Signature (35 genes)
Villus-bottom
Signature (62 genes)
Signature
3
0
5
0
EC
DC1
DC2
EC
DC1
DC2
I
Ly6a
7
0
Avil
5.7
0
J
-Dox
+Dox
CBC
TA
EC
EE
GC
PC
TC
DC1
DC2

### Figure S4

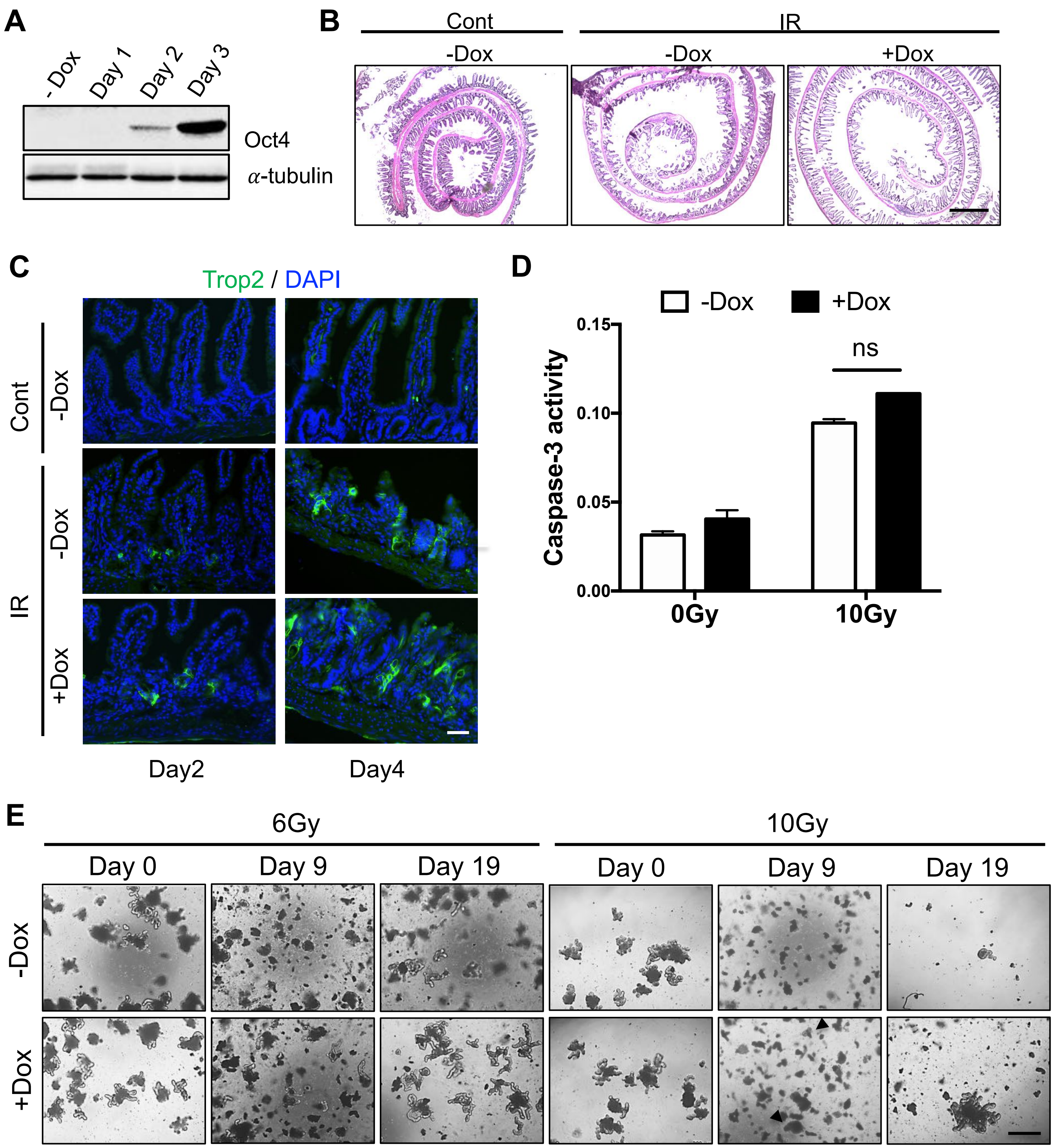
A
- Dox
Day 1
Day 2
Day 3
Oct4
α-tubulin
B
Cont
IR
-Dox
-Dox
+Dox
C
Trop2 / DAPI
Cont
-Dox
IR
-Dox
+Dox
Day2
Day4
D
-Dox
+Dox
0.15
0.10
0.05
0.00
Caspase-3 activity
ns
0Gy
10Gy
E
6Gy
10Gy
Day 0
Day 9
Day 19
Day 0
Day 9
Day 19
-Dox
+Dox

### Figure S5

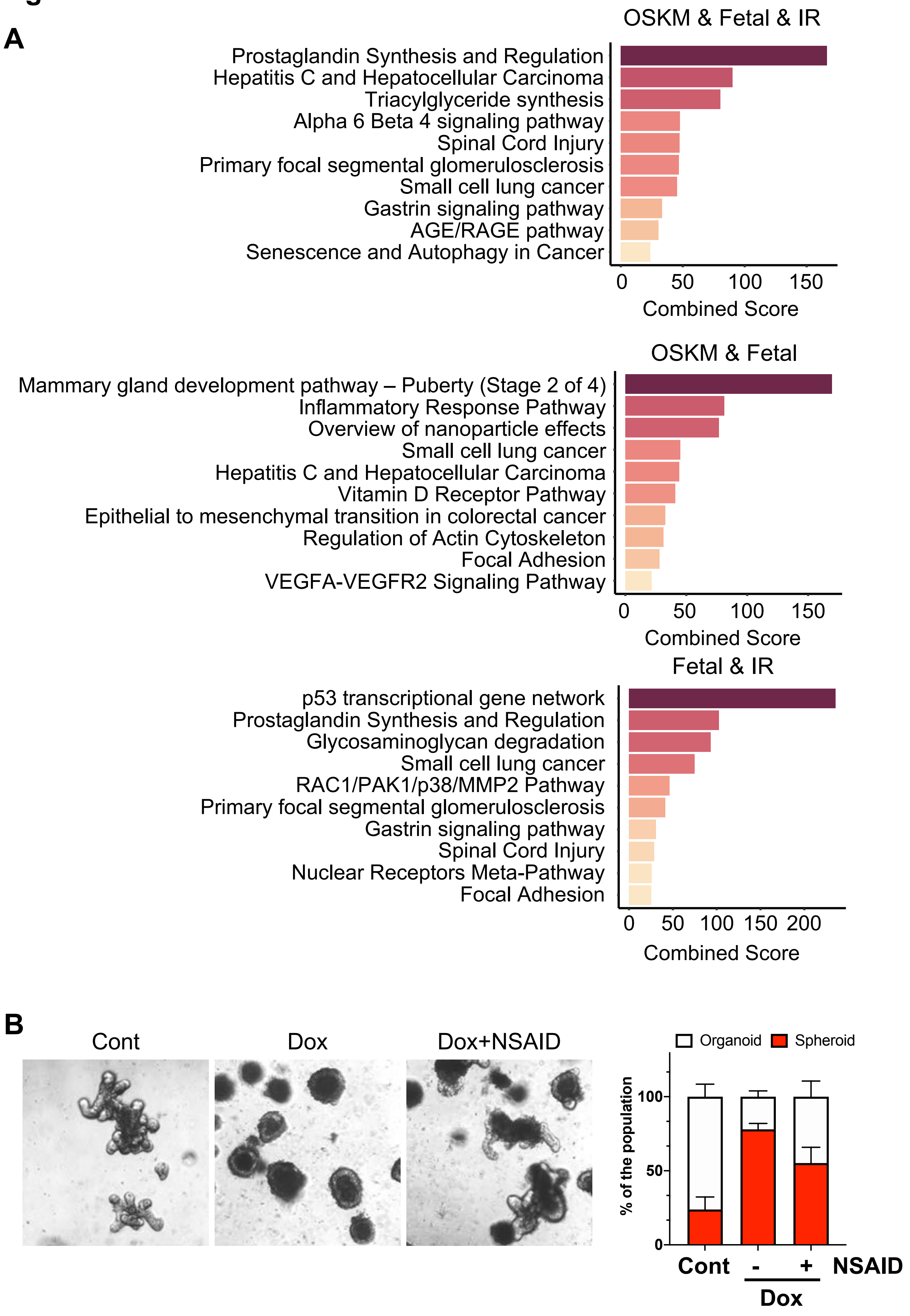
A
OSKM & Fetal & IR
Prostaglandin Synthesis and Regulation
Hepatitis C and Hepatocellular Carcinoma
Triacylglyceride synthesis
Alpha 6 Beta 4 signaling pathway
Spinal Cord Injury
Primary focal segmental glomerulosclerosis
Small cell lung cancer
Gastrin signaling pathway
AGE/RAGE pathway
Senescence and Autophagy in Cancer
0 50 100 150
Combined Score
OSKM & Fetal
Mammary gland development pathway – Puberty (Stage 2 of 4)
Inflammatory Response Pathway
Overview of nanoparticle effects
Small cell lung cancer
Hepatitis C and Hepatocellular Carcinoma
Vitamin D Receptor Pathway
Epithelial to mesenchymal transition in colorectal cancer
Regulation of Actin Cytoskeleton
Focal Adhesion
VEGFA-VEGFR2 Signaling Pathway
0 50 100 150
Combined Score
Fetal & IR
p53 transcriptional gene network
Prostaglandin Synthesis and Regulation
Glycosaminoglycan degradation
Small cell lung cancer
RAC1/PAK1/p38/MMP2 Pathway
Primary focal segmental glomerulosclerosis
Gastrin signaling pathway
Spinal Cord Injury
Nuclear Receptors Meta-Pathway
Focal Adhesion
0 50 100 150 200
Combined Score
B
Cont
Dox
Dox+NSAID
Organoid
Spheroid
% of the population
0
50
100
Cont
-
+
NSAID
Dox

### Figure S5

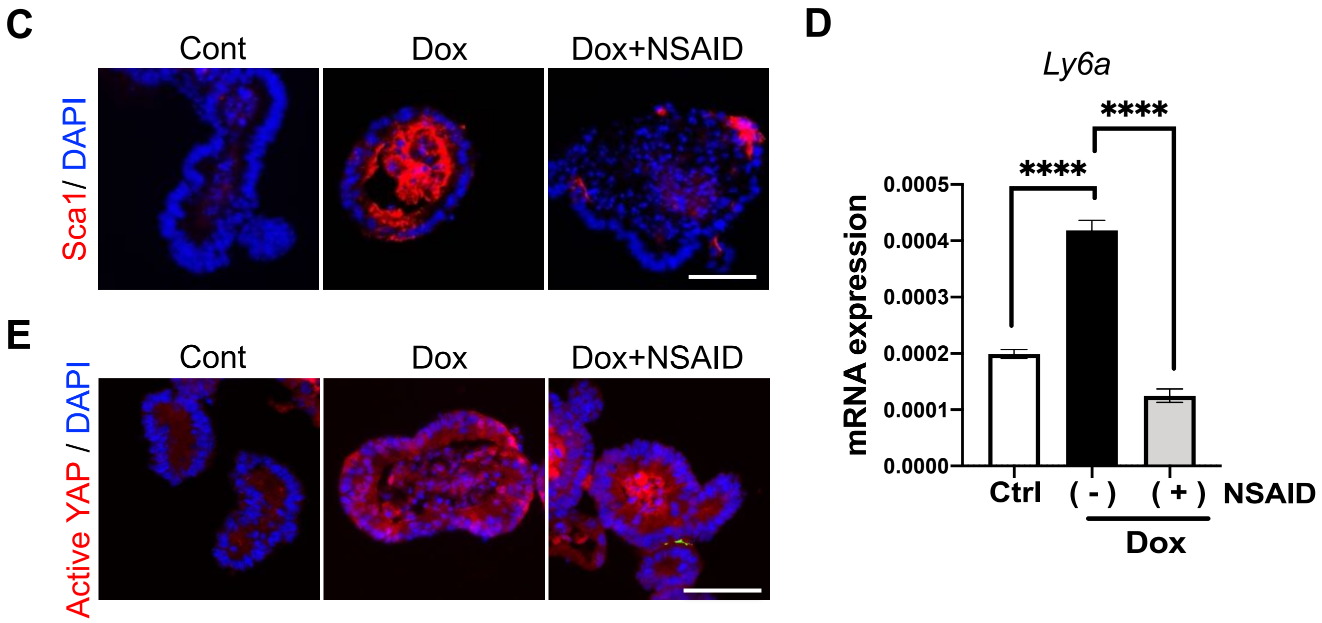
C
Cont
Dox
Dox+NSAID
Sca1/ DAPI
D
Ly6a
****
****
0.0005
0.0004
0.0003
0.0002
0.0001
0.0000
mRNA expression
Ctrl
( - )
( + )
NSAID
Dox
E
Cont
Dox
Dox+NSAID
Active YAP / DAPI

**Figure S6.**

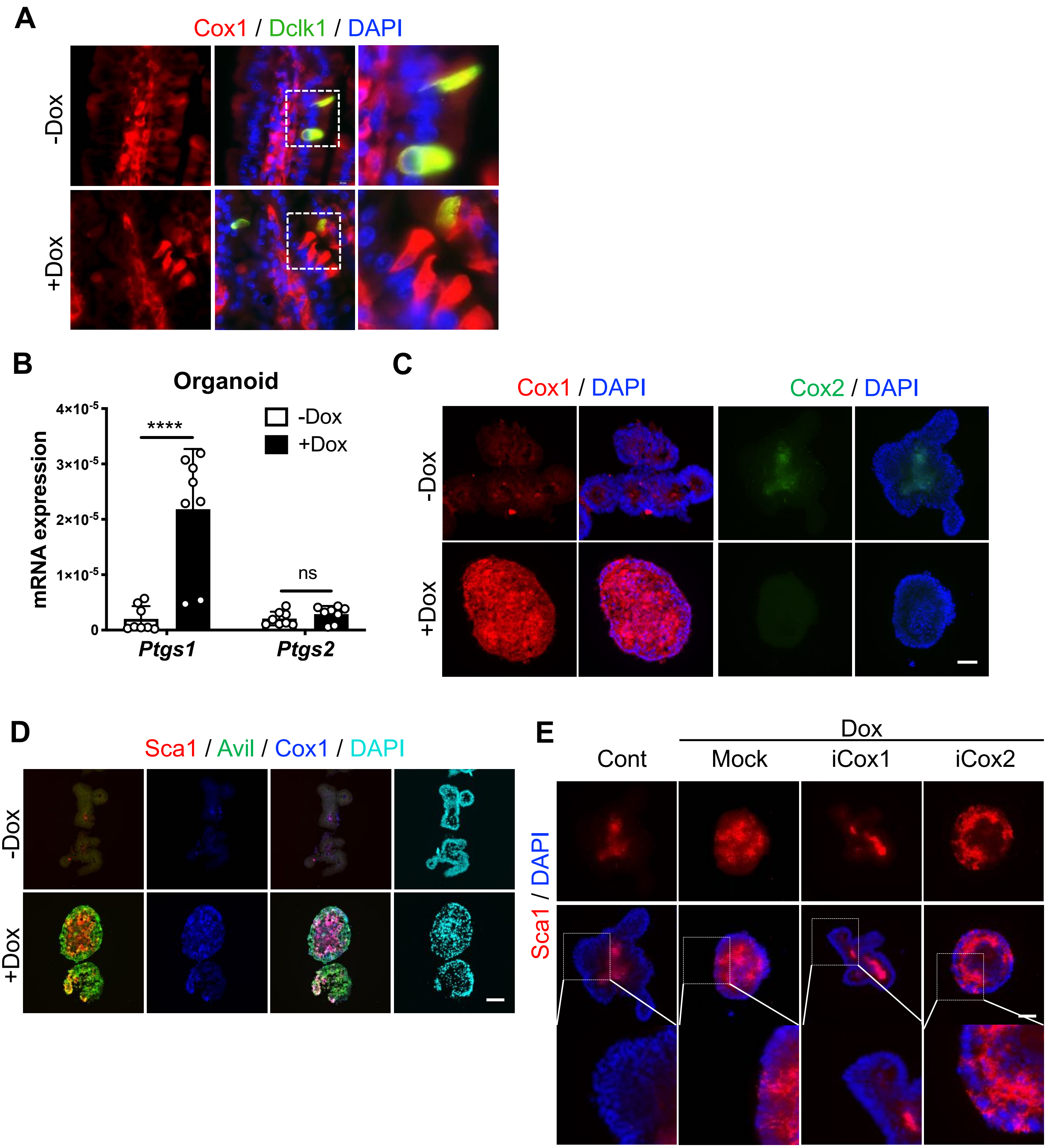
A
Cox1 / Dclk1 / DAPI
-Dox
+Dox
B
Organoid
mRNA expression
4×10-5
3×10-5
2×10-5
1×10-5
0
****
ns
-Dox
+Dox
Ptgs1
Ptgs2
C
Cox1 / DAPI
Cox2 / DAPI
-Dox
+Dox
D
Sca1 / Avil / Cox1 / DAPI
-Dox
+Dox
E
Dox
Cont
Mock
iCox1
iCox2
Sca1 / DAPI
